# Robustness through variability: ion channel isoform diversity safeguards neuronal excitability

**DOI:** 10.1101/2025.09.16.676508

**Authors:** S Hilgert, S Hannah, N Niemeyer, L Huthmacher, S Hürkey, JH Schleimer, S Schreiber, C Duch, S Ryglewski

**Author notes:** shared last and corresponding authorship (JH Schleimer, S Schreiber, C Duch, S Ryglewski). shared first authorship (S Hilgert, S Hannah).

## Abstract

Neural circuits are composed of different neuron types that exhibit distinctly different computational properties resulting from the sets of ion channels expressed. Profound insight exists into how neural computations arise from the precise regulation of ion channels (Armstrong et al., 1998; Lai, Jan, 2006; Nusser et al., 2012), how degenerate channel properties support similar computations (Marder, Prinz, 2002; Marder, Goaillard, 2006), and how channelopathies affect brain function (Kullmann, 2010). However, it remains elusive why neurons express many more channels, and isoforms thereof, than required to tune their specific excitabilities. Here, we employ an experiment-theory approach pairing electrophysiology with *Drosophila* genetics, and mathematical modelling to show that the variance in membrane properties that results from ion channel diversity promotes the robustness of neuron-type specific functions. Specifically, we show that the robustness of flight motoneuron coding properties to internal and external perturbations is significantly enhanced by the diversity of calcium channel splice isoforms expressed. Importantly, increased excitability robustness to perturbations of outward currents or temperature does not require adjustments in calcium channel mean properties. Instead, increases of the variance of calcium channel gating properties that result from channel isoform diversity broaden the dynamic input range the neuron can compute without reaching depolarization block. This broadens our concept of the functional consequences of the tremendous variety and diversity of ion channels expressed in brains.

**One Sentence Summary:** The variance of calcium channel gating properties is increased by channel isoform diversity and aids neuronal coding and excitability robustness.

## Main text

The specific input/output computations that are performed by different types of neurons are key to information processing in brains. Distinct, neuron-type specific excitabilities are implemented by the combinations of ion channels expressed in the neuronal membrane and must robustly be maintained throughout life, despite protein turnover, aging, transcriptional noise, and countless other perturbations. Moreover, the same neuronal excitability class can be encoded by multiple different assortments of ion channels (Prinz et al., 2003; Marder, Goaillard, 2006), thus multiple solutions exist to achieve neuron-type specific target excitabilities. A large body of literature shows that long-term maintenance of neuronal excitability profiles is safeguarded by homeostatic mechanisms that adjust the number and types of ion channel proteins expressed (Marder, 2011; Niemeyer et al., 2021; Wen, Turrigiano, 2024). Much less is known about instantaneous, built-in mechanisms that aid excitability robustness by virtue of the degenerate biophysical properties of the ion channels expressed (Marom, Marder, 2023). Degeneracy, a well described characteristic of the genetic code and the immune system, is defined as multiple structurally different components yielding similar functional output (Edelman, Gally, 2001). Degeneracy in ion channel biophysical properties is an inevitable consequence of the sheer number of ion channel isoforms expressed in the nervous system, e.g., the human genome contains ∼120 genes for voltage gated channels that give rise to multiple 100 to >1000 isoforms with partially overlapping properties. This ion channel degeneracy has been suspected to constitute a neuron intrinsic mechanism to safeguard excitability (Drion et al., 2015; Goaillard, Marder 2021; Albantakis et al., 2024). In contrast to homeostatic mechanisms that actively adjust ion channel densities to tune neurons towards a target region in excitability space, theoretical work suggests that ion channel degeneracy increases the size of that target region (Drion et al., 2015; Goaillard, Marder 2021; Mittal, Narayanan, 2022; Schneider et al., 2023). Here we extend this concept by showing *in vivo* and *in silico* that calcium channel splice isoform diversity renders excitability robust by increasing the variability of the resulting calcium current activation voltage without affecting mean current properties.

### The demands to Drosophila flight MN excitability

We identify and analyze degenerate properties of ion channels that protect neuronal excitability profiles in a highly robust motor circuit that has been optimized over nearly 300 million years of evolution and is used by ∼ 600,000 insect species, the well characterized central pattern generating network (CPG) for asynchronous flight (Hürkey et al., 2023). The circuit consists of 5 motoneurons (MN1-5) that are interconnected by electrical synapses to generate patterned motor output to the “asynchronous” dorsal longitudinal wing depressor muscle (DLM). As described in depth elsewhere (Harcombe, Wyman, 1977; Josephson et al., 2000; Gordon, Dickinson, 2006), in asynchronous flight, motoneuron (MN) action potentials are not in synchrony with muscle contractions, but the MNs fire only every 20^th^ to 40^th^ wingbeat. Importantly, at any given time of flight, all 5 MNs to the DLM fire at the same frequency, and this shared MN firing frequency provides the rate code for how much power output the wing will produce (Gordon, Dickinson, 2006; Hürkey et al., 2023, see Fig. 1a for an example of the relationship between MN firing rate and wingbeat frequency). The input-output computations of the MNs are well-tuned to translate the amount of excitatory input into firing rate and can be quantified by *in situ* current clamp recordings (Fig. 1b). Plotting response firing frequencies as a function of increasing input current amplitudes (F-I curve) reveals a nearly linear translation of input strength into firing rate throughout the working range of MN firing frequencies that are normally used during flight to control wing power (Fig. 1c left, red box depicts normal working range). Importantly, the slope of the F-I curve is smooth and shallow, meaning the MNs encode changes in synaptic drive by small changes in slow tonic response firing rates between 1 and 15 Hz. This has been referred to as class-I excitability (Hodgkin, 1948; Connor, 1975; Ermentrout 1996; Drion et al., 2015), and is an essential feature of MN membrane properties to adequately enable smooth control of wing power during asynchronous flight (Hürkey et al., 2023).

**Figure 1.**
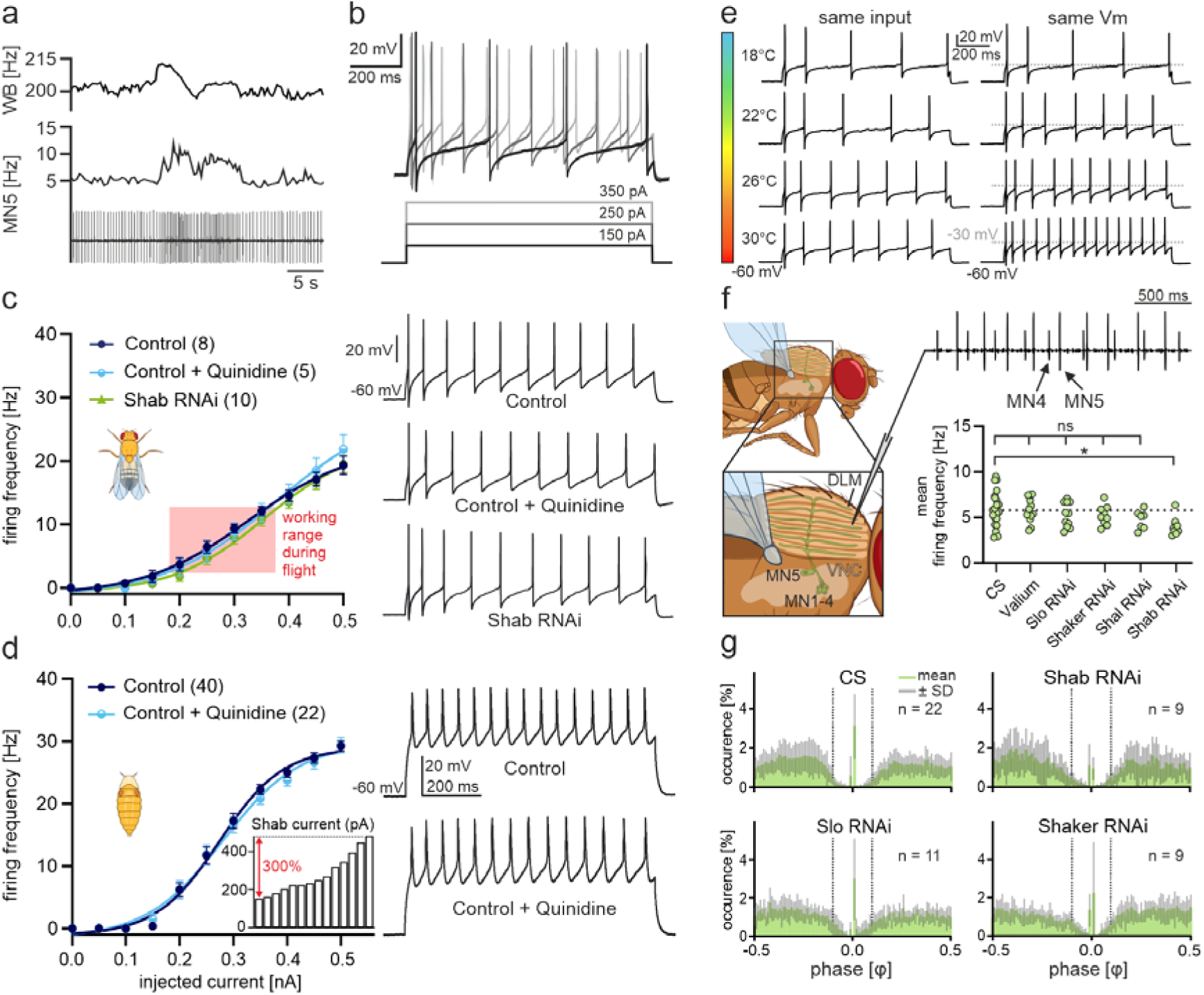
Flight MN excitability and CPG output are robust against perturbations. **a**, MN firing rate (bottom) correlates with and regulates (Hürkey et al., 2023) wingbeat frequency (top). **b**, MN5 displays tonic firing responses to current injection of different amplitudes. **c**, F-I curves (left) reveal class-I MN excitability in controls (dark blue, n = 8) that is robust against genetic (green, n = 10) and acute (light blue, n = 5) blockade of Shab across normal in-flight firing frequency ranges (red box). Example traces on the right. **d**, pupal control MNs (dark blue, n = 40) also exhibit class-I excitability that is robust against Shab blockade (light blue, n = 22). The normal variability of Shab current amplitudes amounts to 300% (inset). **e**, Representative traces show that MN5 retains tonic firing responses to input upon acute temperature manipulation (18° C to 30° C), though response frequencies to the same input (left) or at the same membrane potential (right) are increased. **f**, Schematic of the recording configuration during flight (left) and representative extracellular recording of MN4/MN5 (top right). Mean MN firing frequencies are similar across wildtype (n = 22), a genetic control (Valium, n = 13), *Slowpoke^RNAi^* (n = 11), *Shaker^RNAi^* (n = 9), *Shal^RNAi^* (n = 7) (ANOVA p = 0.36), but *Shab^RNAi^* causes a small but significant reduction (n = 9; Dunnett’s test, p = 0.032). **g**, MN4/MN5 phase relationships are robust against RNAi for different potassium channels. Data as mean ± SEM (c, d) or mean ± SD.

### Drosophila flight MNs display highly robust class-I excitability profiles

According to its key role in flight power control, we predicted MN class-I excitability to be highly robust against internal (ion channel expression levels) and external (e.g. temperature) perturbations. First, we tested the effects of manipulating the K_v_2 potassium channel, Shab, that normally makes up ∼30-50% of the voltage dependent delayed rectifier current in adult and developing flight MNs (see Extended Data Fig. 1). *Shab^RNAi^* with ∼80% knock down efficacy (Extended Data Fig. 2) was targeted selectively to MN1-5 and effective pharmacological blockade of Shab was achieved by acute bath application of 100 µM quinidine (Extended Data Fig. 1). Neither permanent genetic nor acute pharmacological knock down of Shab affect MN tonic firing responses (Fig. 1c, right traces) or MN F-I curves (Fig. 1c, left). This is surprising because Shab is important in flight MNs to control spike onset and phase response dynamics. Both are altered upon overexpression of Shab and significantly affect CPG output (Hürkey et al., 2023). By contrast, slow tonic firing and class-I excitability are neither affected by overexpression (Hürkey et al., 2023) nor knock down of Shab (Fig. 1c). This robustness of class-I excitability could either be due to a broad range of permissive Shab expression levels, or be mediated by compensatory adjustments of other ion channels. Homeostatic adjustments require some time and thus cannot explain the absence of acute pharmacological effects on MN excitability (Fig. 1c), unless the compensatory mechanism operates faster than it takes the drug to act. Since we obtain full blockade of Shab channels after 2 minutes (or faster), we judge compensatory regulation unlikely. Therefore, an intrinsically broad range of permissive Shab expression levels is likely the cause for class-I excitability robustness against the tested perturbations. This is in line with the large variation in the expression levels of Shab, non-Shab, and total delayed rectifier potassium channels that we observe across control animals (Extended Data Figs. 1b, 2), without obvious correlation between the expression levels of different delayed rectifier channels (Extended Data Fig. 1f). Although the normal variation of delayed rectifier amplitude in flight MNs is up to 300% across individuals (Extended Data Fig. 1d), the neurons always robustly exhibit tonic firing and class-I excitability.

A potential source for hidden effects of ion channel manipulation are the large and complex dendritic morphologies of adult flight MNs (∼6000 µm dendritic length and ∼4000 dendritic branches, Vonhoff, Duch, 2010; Ryglewski et al. 2017) which may hamper whole cell voltage control during somatic patch clamp recordings (see methods). To circumvent this problem, we repeated the experiments in far less arborized pupal MNs. Recordings at pupal stage P9 confirmed that developing flight MNs exhibit similar excitability profiles as adult MNs and that acute blockade of Shab does not affect the slow tonic firing responses (Fig. 1d, right), or their class-I F-I curve (Fig. 1d, left). Again, a large variance in different delayed rectifier potassium currents including Shab is observed (Extended Data Figs. 1c, e, see also inset in Fig. 1d) without any obvious correlation in expression levels (Extended Data Fig. 1g). In addition, a large variance in A-type potassium currents and in voltage gated calcium currents is observed (Extended Data Fig. 2). Thus, *Drosophila* flight MN firing responses are robust against substantial animal to animal variability in ion channel expression levels.

We also tested the impact of changes in external conditions on the robustness of MN tonic firing. Temperature is known to affect ion channel properties and the membrane excitability of a neuron (Hesse et al., 2022). Across a range of temperatures that normally occur during the day of a fly but allow development and motor behavior (from 18°C to 30°C) MNs retain their tonic firing (Fig. 1e). Taken together, on the level of single MNs, our *in situ* electrophysiological data show that class-I membrane excitability and slow tonic firing are highly robust against variable ion channel expression levels, acute temperature changes, and perturbations of ion channel expression levels. To test whether the output from the CPG, that is comprised of the 5 electrically coupled MNs (Hürkey et al., 2023), is also robust to perturbation, we targeted ion channel knock downs selectively to all 5 MNs and monitored network output *in vivo* in intact behaving animals. As previously reported (Hürkey et al., 2023), three key characteristics of CPG output are out-of-phase firing of MN pairs (Fig. 1f, top, Fig. 1g), similar firing frequencies across MNs (Fig. 1f, top), and mean MN firing rates of ∼5 Hz during restrained flight (Fig. 1f, bottom). All of these key features of the CPG are robust against genetic knock down of the A-type channels Shaker (K_v_1) and Shal (K_v_4), the delayed rectifier Shab (K_v_2), and the calcium dependent BK channel Slowpoke (Slo). Mean firing rate during sustained flight is ∼5 Hz for all manipulations, although a small reduction in firing rate is observed upon *Shab^RNAi^* (Fig. 1f, bottom). The MN phase relationships as quantified by phase diagrams of the MN4/MN5 pairs (Fig. 1g) remain unaltered. They show a characteristic gap around phase 0 that is a strong indicator for normal phase relationships across all MNs, and thus maximized network desynchronization (which we named splay state, Hürkey et al., 2023). Splay state motor patterns are essential for stable flight control (Hürkey et al., 2023) and are not impaired by genetic knock down of different potassium channels (Fig. 1g). In sum, the CPG network is robust against manipulation of ion channel expression levels in its component MNs, which themselves exhibit large variation in normal ion channel expression levels and are robust against acute internal and external perturbations.

### How to test the functional consequences of ion channel degeneracy in vivo?

What are the mechanisms that render membrane excitability robust? Theoretical and computational studies suggest ion channel degeneracy as a potential source of excitability robustness (Drion et al., 2015, Marom, Marder, 2023; Schneider et al., 2023; Albantakis et al., 2024), but *in vivo* evidence is sparse. *In vivo*, a selective reduction of ion channel degeneracy seems nearly impossible. The challenge is to reduce the number of channel isoforms without affecting mean membrane properties. Only knocking out channels one after the other seems trivial because this will change membrane properties step-by-step until the excitability profile of the neuron is lost. To probe the role of ion channel degeneracy for MN excitability robustness, we, alternatively, reduce the number of ion channels with overlapping properties without affecting the average properties of the MN membrane currents. To achieve this, we reduce the splice isoform diversity of the voltage gated calcium channel (VGCC) cacophony (cac, the fly Ca_v_2 homolog).

In *Drosophila melanogaster*, alternative splicing leads to 18 annotated cac channel isoforms. By targeted on-locus CRISPR-Cas9 mediated alternative exon excision, the number of possible cac isoforms can be reduced (see methods and Bell et al., 2025). *Drosophila* flight MNs express cac channels in the somatodendritic domain, along the axon, and in presynaptic active zones in the axon terminal (Kawasaki et al., 2004; Ryglewski et al., 2012; Heinrich, Ryglewski, 2020). Importantly, only specific cac splice isoforms localize to presynaptic active zones and are required for synaptic transmission. By contrast, along the MN central arbors that underlie spike generation and excitability control, cac isoform abundance is less regulated. In fact, cac isoforms required in active zones, as well as isoforms with no synaptic function, show functional expression in MN central arbors (Bell et al., 2025). Therefore, the diversity of cac isoforms with overlapping and thus degenerate functions that contribute to MN excitability control is high.

Excision of the two mutually exclusive exons *IS4A* and *I-IIA* (*ΔIS4A* Δ*I-IIA*) results in a ∼80% reduction of cacophony isoform diversity from 18 to 4 isoforms (Bell et al., 2025). Importantly, this does not change the characteristic slow tonic firing and class-I excitability of adult or of pupal MNs as measured by F-I relations (Fig. 2a, b). Moreover, total calcium current as recorded from the soma of pupal MNs remains mostly unchanged upon reducing cac isoform number from 18 to 4 in *ΔIS4A ΔI-IIA* as compared to controls with full isoform diversity. In controls and in *ΔIS4A ΔI-IIA*, cac currents have qualitatively similar kinetics (Fig. 2c), activate between -60 mV and -50 mV, show a mean half activation at about -25 mV and maximum amplitudes at 0 mV, which results in similar I-V curves (Fig. 2d). However, the I-V curves of *ΔIS4A ΔI-IIA* neurons display a small shift at potentials between -30 mV and +20 mV (Fig. 2d) going along with a small but statistically insignificant increase in the amplitude of I_max_ (Figs. 2d, e; p = 0.5221, unpaired T-test). The resulting difference between the I-V curves with full and with reduced cac channel isoform diversity is small and not reflected in statistically significant differences in mean current amplitudes or activation voltages, but it is plotted as a difference current (I_Ca_, Fig. 2d, red plot) because it is functionally relevant (see below).

**Figure 2.**
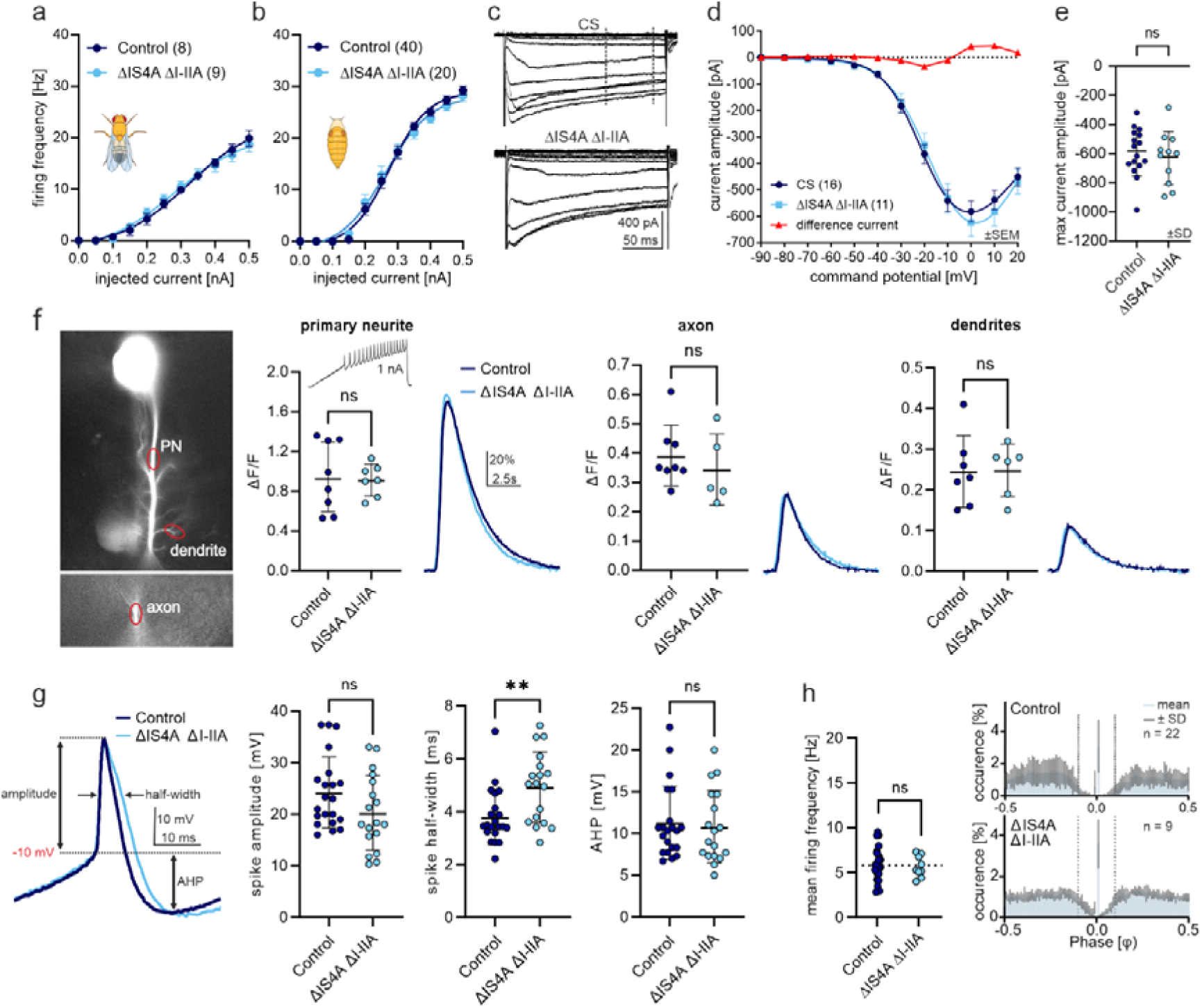
Reducing VGCC isoform diversity does not affect mean current properties. Excision of *IS4A* and *I-IIA* (Δ*IS4A* Δ*I-IIA*) reduces channel isoform number from 18 to 4. This does not affect tonic firing and class-I excitability of adult (**a**) or pupal (**b**) MNs. **c**, whole cell pupal MN calcium currents remain similar with full or reduced isoform numbers. **c, d**, in controls and in Δ*IS4A* Δ*I-IIA*, cac currents have similar kinetics (c), activate between -60 mV and -50 mV (**d**), show a half activation at about -25 mV (**d**) and maximum amplitudes at 0 mV (**d**). Isoform reduction induces a small shift of the I-V curve that is plotted as difference current (I_Ca_ = control-*ΔIS4A ΔI-IIA*) in red (**d**) but is statistically not significant (ANOVA, p = 0.85). **e**, I_max_ is not significantly affected by isoform reduction (p = 0.5221, unpaired T-test). **f**, GCaMP6s imaging of calcium signals upon spike induction by somatic current injection in primary neurite (PN), axon, and dendrites shows no significant differences (unpaired T-test, PN p = 0.8278, axon p = 0.4691, dendrites p = 0.9570). **g**, spike shape analysis reveals no significant differences in maximal spike (Mann-Whitney U-test, p = 0.0981) or AHP amplitude (unpaired T-test, p = 0.7360), while spike half-width in *ΔIS4A ΔI-IIA* is increased (Mann-Whitney test, p = 0.0060). **h**, mean firing frequencies during flight (left) (unpaired T-test, p = 0.6903) and phase relations between MN4/5 pairs (right) are unaffected. Data shown as mean ± SEM (**a, b, d**) or mean ± SD (**e, f, g**).

Calcium imaging reveals no differences in activity dependent calcium influx into the different sub-neuronal compartments of these MNs, primary neurite, axon, or dendrites, upon reducing cac channel isoform number in *ΔIS4A ΔI-IIA* (Fig. 2f). These data are consistent with the interpretation that a reduction of cac channel isoform diversity does not go along with major changes in the mean values of cac current properties. Importantly, a reduction in cac channel isoform number does also not affect Shab mediated, or non-Shab delayed rectifier currents, or total A-type currents (Extended Data Fig. 3). This indicates that the reduction in cac channel isoform diversity does not affect mean values of voltage gated calcium or potassium currents. Similarly, at the level of CPG output during flight, reduced VGCC isoform diversity has no effect on mean MN firing frequencies (Fig. 2h left), and it does not affect the characteristic phase relationships of MN pairs as revealed by phase histogram (Fig. 2h, right). However, we observe an effect on spike shape. Although the excision of the exons *IS4A* and *I-IIA* has no effect on the amplitudes of the spike overshoot or the afterhyperpolarization (AHP), spike half-width is prolonged by ∼25% (Fig. 2g).

Taken together, reducing isoform number of cacophony VGCCs from 18 to 4 by excision of the alternative exons *IS4A* and *I-IIA* does not significantly affect MN input-output computation as determined by F-I curves, spike or AHP amplitude, the mean properties of the somatodendritic calcium current, activity dependent calcium influx, voltage gated potassium currents, or coordinated CPG output. However, small, yet statistically insignificant, shifts in the I-V relationship go along with a functional difference, an increased action potential half width. We conclude that calcium channel isoform number can be reduced *in vivo* without affecting the mean membrane current properties that underlie class-I membrane excitability and can thus be used to test the role of cac channel isoform degeneracy for excitability robustness.

### VGCC isoform diversity protects MN tonic firing from knock down of Shab current

To test the hypothesis that ion channel degeneracy provides neuronal excitability robustness, we next combined our unexpected result that MN class-I F-I curves and MN tonic firing are robust against acute blockade of Shab (Figs. 1c, d) with the ability to selectively manipulate cac channel isoform degeneracy. We specifically tested whether MN sensitivity to pharmacological block of Shab potassium channels is significantly increased in the background of reduced VGCC isoform diversity.

Reducing cac isoform number alone does not affect MN class-I excitability (Figs. 2a, b), and acute blockade of Shab does not alter MN5 firing responses to somatic current injection in the presence of full VGCC isoform diversity (Figs.1c, d). In contrast, with reduced VGCC isoform diversity *(ΔIS4A* and *ΔI-IIA*), the same blockade of Shab causes a range of severe phenotypes in spike shape and slow tonic firing responses (Fig. 3a). We categorized and color-coded the firing phenotypes observed upon somatic current injection into 5 types (Fig. 3a): tonic firing responses with normal spike shapes (Fig. 3a, top trace, white), tonic firing but spike broadening (Fig. 3a, second from top, yellow), double spikes during tonic firing (Fig. 3a, third trace from top, orange), plateau-like spikes (Fig. 3a, second from bottom, red), and depolarization block (Fig. 3a, bottom trace, magenta). In each recorded MN, multiple sweeps of somatic square pulse current injection at increasing amplitudes (see methods) were applied. The different spike shapes exemplified in Fig. 3a can occur in subsets of spikes within a sweep, or in subsets of sweeps, but the phenotypic severity becomes higher for sweeps with higher amplitude current injection (Fig. 3b). We next quantified the occurrence of the different spiking phenotypes across genotypes as the relative amount of time spent spiking in any given spiking category and plotted the data as color coded stacked bar charts for each genotype (Fig. 3c).

**Figure 3.**
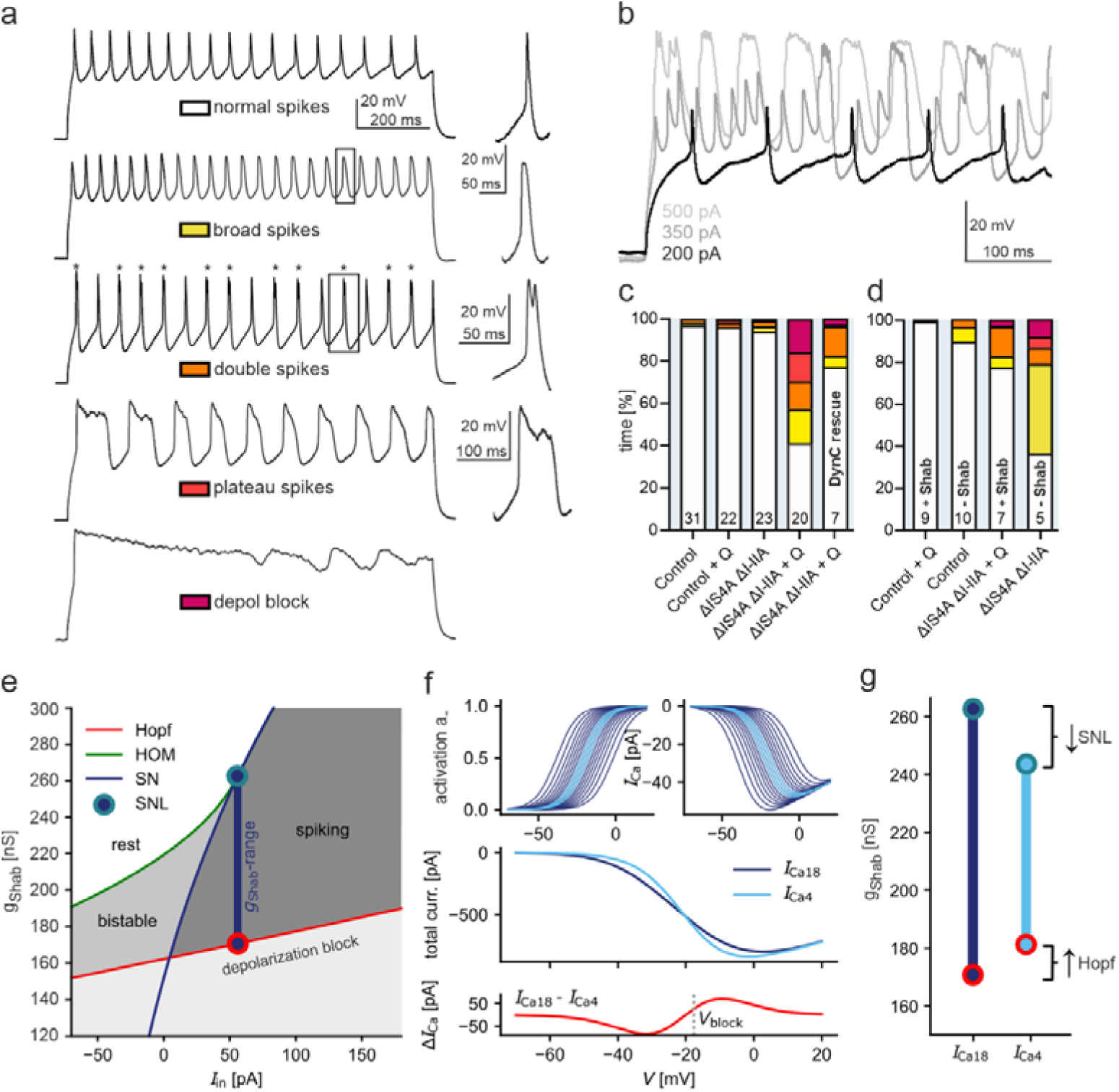
VGCC isoform diversity aids excitability robustness. **a**, Spike shape categories observed upon Shab blockade in MNs with reduced VGCC isoform diversity (*ΔIS4A ΔI-IIA*). The severity of the phenotype does not necessarily correlate with absolute amplitude of current injection and can differ between cells. From top: normal spike shape (white), broadened spikes (yellow), double spikes (orange), plateau spikes (red), depolarization block (magenta). **b**, Phenotypic severity increases with input amplitude c, Quantification as percent time different spiking categories occurred. Quinidine (Q) blockade of Shab (n = 22) alone or *ΔIS4A ΔI-IIA* (n = 23) alone do not impact normal spiking. Shab blockade in *ΔIS4A ΔI-IIA* (n = 20) results in severe spiking phenotypes, but normal MN spiking is partially rescued by restoring Shab with dynamic clamp (n = 7, right bar) **d**, Same (*ΔIS4A ΔI-IIA*) as (**c**), but manipulating Shab with dynamic clamp. **e**, Codim-2 bifurcation diagram showing the permissive Shab range in MN models (vertical blue line). The upper boundary is defined by the SNL point (teal circle) at the shift from the homoclinic into the SNIC excitability class. The Hopf line (red) defines the lower boundary, where MNs approach depolarization block. **f**, VGCC splice isoforms are modeled by varying their half-activation voltages V_½_ (top traces). Full isoform diversity resulted in 18 splice channels with a large spread around the mean V_1/2_ (dark blue) while 4 splice currents spread less around V_½_ (light blue). Steady-state I-V curve shows a shallower slope with 18 (control) than with 4 (*ΔIS4A ΔI-IIA*) splice currents (middle). Subtracting I-V plots with 4 and 18 isoforms yields ΔI_Ca_ (bottom trace, red). **g**, In the model, splice isoform reduction decreases the permissive Shab range.

In all control animals with full VGCC isoform diversity and no Shab channel manipulation (n=31), tonic firing with normal spike shape was observed across all current injection amplitudes, and only in very rare cases a singled-out altered spiking category was observed (broad spikes in 1.38%, and double spikes in 1.62% of the time). The same applies to acute blockade of Shab in animals with full VGCC isoform diversity (n=22, double spikes: 2.16%, plateau spikes: 1.44%, depolarization block: 0.4%) and animals with reduced VGCC isoform diversity (n=23, broad spikes: 2.44%, double spikes: 2.7%, depolarization block: 0.85%). In contrast, upon blockade of Shab in animals with reduced VGCC isoform diversity (n= 20), tonic firing with normal spike shape was observed in only 40% of total time spiking. The remaining 60% of firing responses are composed of broad spikes (16.28%), double spikes (13.16%), plateau spikes (13.81%), and depolarization block (15.69%). Depolarization blocks and plateau-like spikes are the most severe phenotypes and impair MN rate coding. Less severe phenotypes, such as broad and double spikes, decrease the fidelity of rate coding. These data show that the normal robustness of MN tonic rate coding to perturbation of Shab outward current is significantly reduced by a reduction in the numbers of VGCC isoforms (p < 0.0001, Fisher’s Exact test), or *vice versa*, normally, the expression of degenerate VGCC isoforms renders tonic firing robust. Finally, the significant effects of Shab block in MNs with reduced VGCC isoform diversity can be partially rescued by restoring Shab current in the recorded MN by dynamic clamp (Fig. 3c, right bar). Mild spiking phenotypes (broad and double spikes) are fully rescued and severe spiking phenotypes (plateaus and depolarization block) are transformed into mild spiking phenotypes (Fig. 3c, right bar). To exclude unwanted side effects of quinidine and to restrict the Shab manipulation only to the recorded neuron, we also compared knock out of Shab current by dynamic clamp (Fig. 3d). First, in controls with full VGCC isoform diversity, slow tonic firing is unaffected by acute pharmacological or dynamic clamp block of Shab (Fig. 3d, left two bars). Second, with reduced VGCC isoform diversity, robustness of tonic firing declines not only upon pharmacological but also upon dynamic clamp block of Shab (Fig. 3d, right bar). Importantly, the few instances when aberrant firing phenotypes are observed upon Shab manipulation in neurons with full VGCC isoform diversity occur at the upper limit or beyond the MN firing frequencies that are observed during normal behavior. By contrast, with reduced VGCC isoform diversity aberrant firing phenotypes are induced by Shab manipulation within the firing rate range that is behaviorally relevant and observed during flight (see Extended Data Fig. 4). Taken together, VGCC isoform diversity provides a MN intrinsic mechanism that renders tonic firing robust against acute ion channel manipulation.

Therefore, our experimental data are consistent with a role of ion channel degeneracy in neuronal excitability robustness. Specifically, without changing the mean properties of calcium currents or any other current measured from these MNs, the reduction of VGCC isoform number increases excitability vulnerability to potassium channel manipulation (Fig. 3). However, given the complexity of the dendritic structure and the ion channel complement of these MNs, that we did not measure all membrane currents, and that we observe a small though statistically insignificant shift of the I-V curve (Fig. 2d) and an effect on action potential shape (Fig. 2g), our experiments cannot exclude the possibility that the removal of the *IS4A* and *I-IIA* exons from the *cacophony* gene affects membrane properties that are hidden to our tests and that affect MN excitability robustness. Therefore, we next used computational models that allow keeping all other parameters constant, to then test whether channel isoform diversity alone is sufficient to increase excitability robustness.

### Bifurcation analysis reveals effects of VGCC isoform diversity on excitability robustness

We previously provided mathematical, modeling, and experimental evidence that correct motor output from the flight CPG requires the MNs to exhibit (i) class-I excitability to control flight power output by rate coding and (ii) a homoclinic spike onset that favors network desynchronization and thus stable flight (Hürkey et al., 2023). From a dynamical systems theory perspective (see methods for details), these requirements define an upper limit of Shab expression (Hürkey et al., 2023) that can be identified from the bifurcation diagram (Fig. 3e): Above the so-called saddle node loop (SNL) point, MN excitability shifts from the homoclinic into the SNIC regime (Fig. 3e). We previously showed that further into the SNIC regime, the CPG network loses normal motor coordination, namely the splay state pattern (see Hürkey et al., 2023). The new data presented here on single MN spike waveforms define a lower level of tolerable Shab expression, because Shab reduction can cause plateau potentials and depolarization block (Fig. 3a). Within the bifurcation diagram, these features can be observed close to and below the Hopf line (see methods) where MN firing responses approach depolarization block (Fig. 3e), thus predicting aberrant spiking phenotypes and a loss of rate coding as observed experimentally (Figs. 3a-d). Therefore, from a dynamical systems theory point of view, we define a tolerable Shab range (Fig. 3e, vertical blue line) with an upper (Fig. 3e, SNL point) and a lower limit (Fig. 3e, Hopf line) of Shab channel levels in codim-2 bifurcation diagrams. This region defines the Shab range in which single MN tonic firing, as well as the network coordination (splay state) are guaranteed.

To test by further bifurcation analyses whether VGCC isoform diversity can affect the permissive Shab range, we extended our previously validated three-dimensional model of *Drosophila* flight MNs (Hürkey et al., 2023) with cac VGCCs based on *in situ* recordings. Since we know that different cac splice isoforms that are expressed in the somatodendritic domain of *Drosophila* flight MNs differ in their half activation voltages (*V_½_*, Bell et al., 2025), but the mean half activation voltage remains unchanged upon excision of the exons *IS4A* and *I-IIA* (Fig. 2d), we implemented different degrees of cac channel isoform diversity by varying the isoform *V_½_*. We further assumed that all VGCC isoforms are expressed in equal proportion and that the slope of their activation curve is identical. Full isoform diversity is accounted for by 18 splice currents with different *V_½_* that spread at equal increments around the mean (Fig. 3f, top panels, dark blue). Reduced isoform diversity is accounted for by 4 splice currents (modeling excision of *IS4A* and *I-IIA*) that spread less around the mean *V_½_*(Fig. 3f, top panels, light blue). The rationale for choosing a smaller spread around the mean after excision of *IS4A* and *I-IIA* is the high sequence similarity of the remaining 4 splice isoforms (see flybase.org). In sum, our model restricts the difference between full and reduced VGCC isoform diversity to the variance of *V_½_*.

When comparing the permissive Shab range of models with full and reduced VGCC isoform diversity, it becomes evident that the range in which MNs display tonic firing and have a phase susceptibility that favors network desynchronization decreases with reduced VGCC isoform diversity (Fig. 3g). This observation is consistent with our experimental data and the idea that the variance of VGCC gating properties aids excitability robustness. We next used this model to further investigate the underlying mechanism, and to derive experimentally testable model predictions.

### A mechanism for excitability robustness through the variance of VGCC gating properties

Bifurcation analysis reveals that reducing isoform diversity shrinks the permissive Shab range of MNs by moving the depolarization block (Hopf line) and the SNL switch closer together (Figs. 3e, g). Both of these shifts can be explained when comparing the steady-state I-V curves (Fig. 3f, middle) and the dynamic calcium currents (Fig. 4a, Extended Data Fig. 5a) during a spike of models with full and reduced VGCC isoform diversity. Due to the larger variance of *V_½_* in the presence of all isoforms, the effective I-V curve is, in sum, shallower than in the splice-reduced scenario. Subtraction of the summed calcium current with reduced isoform diversity from that with full isoform diversity defines a difference-current, Δ*Ι*_Ca_ = *I*_Ca18_-*I*_Ca4_ (Fig. 3f, bottom, red trace). Δ*Ι*_Ca_ arises from the increased variance in *V_½_* of the additional splice isoforms in the wildtype and offers an explanation for the experimentally observed increased excitability robustness. Notably, a similar Δ*Ι*_Ca_ is seen experimentally when subtracting the control cac current I-V curve from the one obtained with excision of *IS4A* and *I-IIA* (see Fig. 2d). The immediate observation is that, for models with full isoform diversity, Δ*Ι*_Ca_ provides a repolarizing force at elevated membrane voltages around the depolarization block and a depolarizing force at more negative voltages that are covered during the refractory period. In fact, the voltages at which the depolarization block occurs *in vivo* coincide with membrane voltages at which the difference current, Δ*Ι*_Ca_, provides the reduced depolarization (see Fig. 2d). Therefore, the difference current that results solely from different degrees of *V_½_*variance could in principle prevent the spiking phenotypes that are observed *in vivo* with reduced VGCCs isoform diversity and reduced Shab current, i.e. spike broadening, double spikes, plateau potentials, and depolarization block (see Figs. 3a-d).

**Figure 4.**
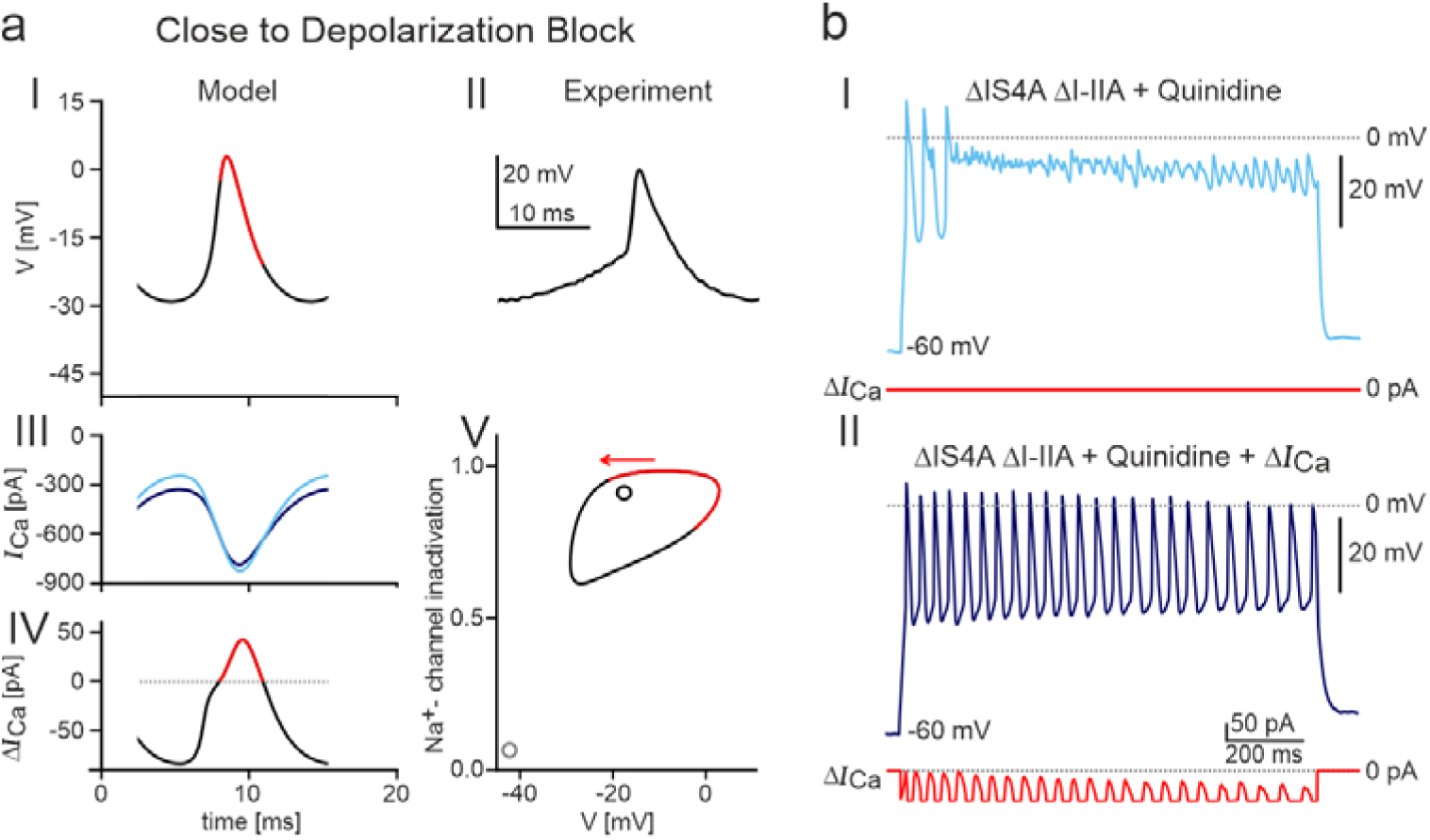
VGCC isoform diversity protects from depolarization block. **a**, Simulations of membrane voltage and calcium current during an action potential close to the depolarization block. First row shows model voltage traces during an action potential (**I**) that are similar to those recorded in vivo (**II**). Second row depicts the dynamic calcium currents during an action potential for models with 4 (light blue) and 18 (dark blue) VGCC isoforms (**III**). In the third row (**IV**), the differences of the dynamic currents (dynamic Δ/_Ca_) are shown over the time course of the action potential. **V**, phase portrait of an action potential. The ghost of the saddle-node (i.e., old resting state) is shown as a grey open circle. An unstable spiral (i.e., location of the depolarization block) is indicated by a black open circle. The reduced depolarization during action potential downstroke (red) destabilizes the depolarization block and supports tonic spiking. **b I & II**, representative example of the firing response of a MN with reduced VGCC isoform diversity and Shab blockade by quinidine close to depolarization block **(I)**. Adding Δ*Ι*_Ca_ with dynamic clamp to the same recording rescues the tonic firing response **(II)**.

A closer look at the difference in dynamic calcium currents between models with full and reduced VGCC isoform diversity during MN spiking reveals how the splice current both destabilizes the depolarization block (Fig. 4a) and stabilizes the SNL point (Extended Data Fig. 5a) and thus increases the robustness range. *In vivo* and *in silico,* spikes decrease their amplitude when approaching the depolarization block (compare Fig. 4a I & II and Extended Data Fig. 5a I & II). The dynamic calcium currents during spiking display clear differences in models with 4 (light blue) versus 18 (dark blue) calcium channel isoforms (Figs. 4a III, Extended Data Fig. 5a III). To isolate these differences, we examine the dynamic difference current (Fig. 4a IV and Extended Data Fig. 5a IV). The differential impact of Δ*I*_Ca_ during different phases of the action potential provides a mechanistic explanation for why the permissive Shab range decreases with reduced VGCC isoform diversity.

During action potential peak and early downstroke, models with full isoform diversity generate less depolarizing calcium current (Δ*I*_Ca_ > 0). Simulations show that this reduction in depolarization (Δ*I*_Ca_ > 0, Figs. 4a I, IV & V, red) prevents the neuron from entering the depolarization block. Conversely, models with reduced VGCC isoform diversity experience a stronger depolarization. As a result, the lower bound of the functional Shab range shifts upward, since a greater hyperpolarizing force, through increased Shab conductance, is required to sustain regular spiking (Figs. 3e, g).

Our simulations show that, mechanistically, an increased variance of VGCC gating properties can cause a small reduction in the calcium current amplitudes at precisely those membrane voltages where increased input causes depolarization block. To test experimentally whether Δ*Ι*_Ca_ as resulting from differences in variance of *V_½_*can effectively prevent depolarization block in MNs, we use dynamic clamp to add Δ*Ι*_Ca_ to a current input that causes aberrant spiking phenotypes in a *ΔIS4A ΔI-IIA* MN with pharmacological Shab block (Fig. 4b I). We approximated Δ*Ι*_Ca_ using an analytical derivation (see methods) that was then fit to I-V plots of control and *ΔIS4A ΔI-IIA* MNs (Fig. 2d). Adding Δ*Ι*_Ca_ indeed abolishes the depolarization block and results in a tonic firing response in the same recording (Fig. 4b I & II). These data further support our interpretation that the variance of VGCC gating properties safeguards MN excitability profiles.

At the upper boundary of the permissive Shabe range, marked by the SNL bifurcation, models with reduced VGCC isoform diversity show a reduction of the boundary to lower conductances of *g_Shab_* (Fig. 3g), thereby further decreasing the range. As mentioned above, the SNL marks a switch in a neuron’s excitability class from the homoclinic to the SNIC regime which impacts the network synchronization state, yet only subtly affects spike phenomenology. The excitability switch is associated with changes in the action potential trajectory in the phase portrait (Extended Data Fig.5a V). Whereas neurons of the homoclinic class approach the saddle point along the strongly stable manifold, SNIC neurons approach along the slow center manifold. The increased depolarization during late action potential downstroke and refractory period with more VGCC isoform diversity (Δ*I*_Ca_ < 0, Extended Data Fig. 5a, I, IV & V, teal) essentially prevents SNIC dynamics and thus maintains the homoclinic spike onset (Extended Data Fig. 5 V, teal arrow). As the excitability switch at the SNL is linked to changes in network synchronization (Hürkey et al., 2023), not measurable in single cell patch-clamp recordings, we next performed network simulations to test whether full VGCC isoform diversity expands the range in which the network desynchronizes. The simulations reveal that networks with full VGCC isoform diversity can sustain desynchronization across a larger parameter range (Extended Data Fig. 5b, 5c). Taken together, the particular form of Δ*Ι*_Ca_ increases the *Shab* range by differentially repolarizing near the depolarization block and depolarizing near the SNL point. Sensitivity analysis reveals that changing model dynamics (e.g., by changing *V_1/2_*) affects this differential polarization and therefore also the impact on the Shab range (Extended Data Fig. 6).

### VGCC isoform diversity protects MN tonic firing across different temperatures

Does this robustness through VGCC isoform diversity hold across different external conditions, such as temperature changes within the range that a fly normally faces (from 18°C to 30°C)? And how do temperature and the permissive Shab range interact? Since it is well established that temperature directly affects neuronal excitability (Hesse et al., 2022), in poikilotherm animals, mechanisms to protect neuronal input/output computations must operate over the full range of physiological temperatures. We already know that normal MN slow tonic firing is stable across this temperature range (see Fig. 1e). Can our models predict the effects of temperature on the permissive Shab range and excitability of MNs with full or reduced VGCC isoform diversity? To investigate this, we introduced temperature-dependent kinetics to our model (see methods for detail) and constructed temperature-dependent codim-2 bifurcation diagrams for MNs with full isoform diversity (Fig. 5a) and for Δ*IS4A* Δ*I-IIA* MNs with reduced isoform diversity (Fig. 5b). We further tested our model prediction by *in vivo* recordings with adjustable bath temperatures (Figs. 5d, e). Since control MNs are normally robust across the 18°C to 30°C temperature range (see Fig. 1e), we acutely blocked Shab by bath application of quinidine and replicated this manipulation in the model by reducing *g_Shab_* by 10% from its value at the SNL. As previously shown for other neurons (Hesse et al., 2022), increases in temperature shift excitability deeper into the HOM space toward the Hopf line and depolarization block (Figs. 5a, b). At 22°C, MNs with full VGCC isoform diversity retain tonic firing responses to current injection in the face of Shab block (Figs. 1c, d). As predicted by the model (Fig. 5a), temperature increases to 30° C shift the neuron more towards the depolarization block into a region where aberrant spiking phenotypes (see Fig. 3a) can be evoked. The phenotypes can be fully reverted by lowering the bath temperature again from 30°C to 22° C in the same recording (Fig. 5d). Therefore, acute temperature changes can be used to shift excitability back and forth (Fig. 5d) close to the Hopf line (Fig. 5a). Further, comparing the bifurcation diagram of models with full (Fig. 5a) and reduced (Fig. 5b) VGCC isoform diversity reveals that lowering isoform diversity moves the SNL and depolarization block closer together. These shifts thereby decrease the temperature region in which model neurons maintain tonic firing and have a phase susceptibility that favors desynchronization. This model prediction is also confirmed in experiments, where in MNs with reduced VGCC isoform diversity, bath application of quinidine causes aberrant spiking already at 22° C (Fig. 5e). Given that the bifurcation diagrams for Shab (Fig. 3e) and temperature (Fig. 5a, b) are nearly mirror images of each other, we reasoned that altering one parameter (e.g., blocking Shab) could be compensated by adjusting the other (e.g., decreasing temperature). This prediction is confirmed *in vivo*, because acutely lowering the bath temperature from 22° C to 18° C restores tonic firing, and raising it to 22° C again causes aberrant firing (Fig. 5e).

**Figure 5.**
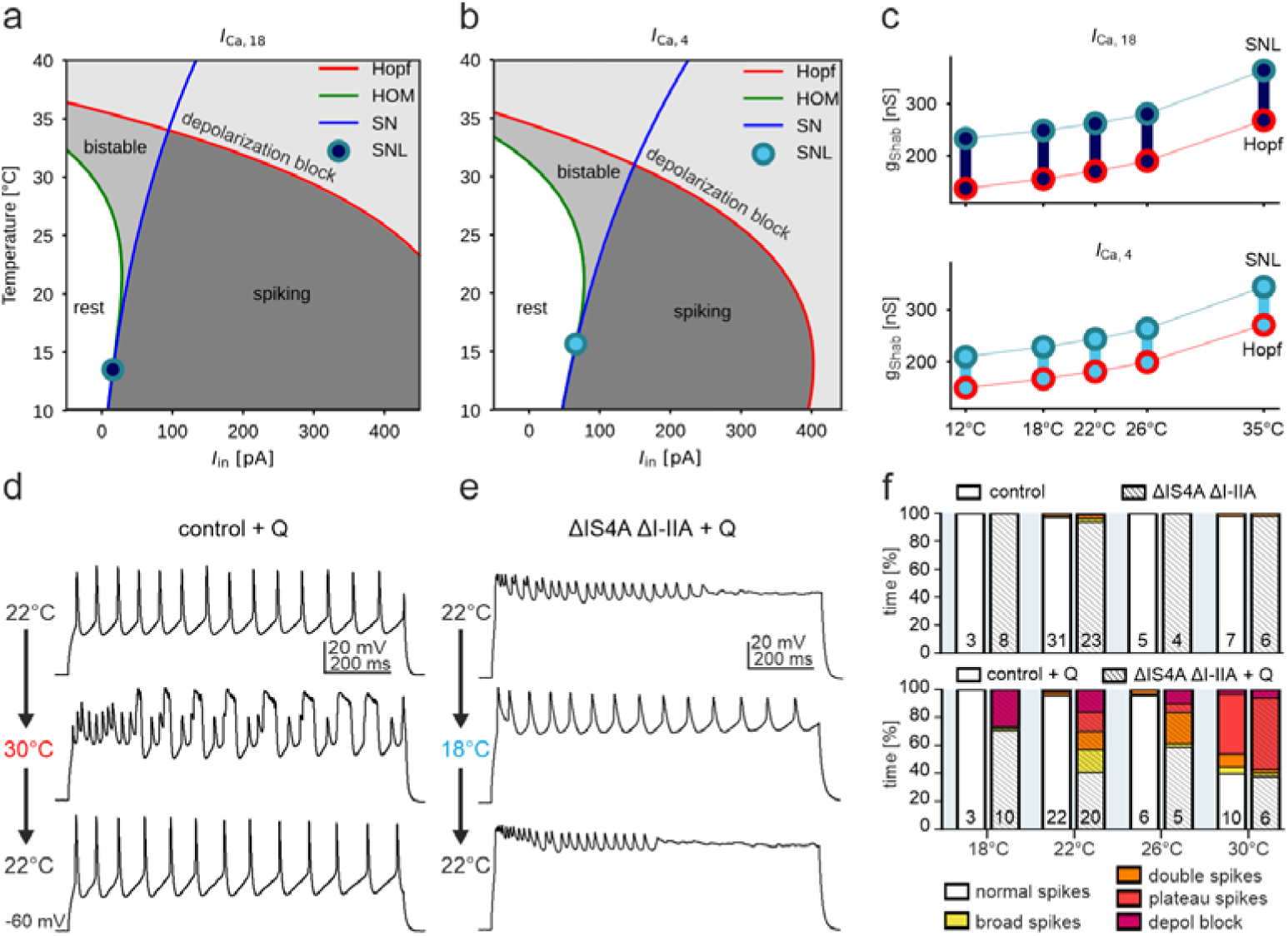
VGCC isoform diversity aids excitability robustness against temperature. Temperature-dependent codim-2 bifurcation diagram for models with (**a**) full (*I*_Ca,18_) and **(b)** reduced (*I*_Ca_, _4_) VGCC isoform diversity after reducing *g*_Shab_ by 10% to simulate quinidine manipulation. **c**, functional Shab range at different temperatures between 12°C and 35°C for models with full (dark blue) and reduced (light blue) isoform diversity. **d**, firing responses to the identical somatic current injection of a MN with full VGCC isoform diversity and acute Shab blockade (+Q) at 22°C (top trace), 30°C (middle) and back at 22°C (bottom). **e**, firing responses to the identical somatic current injection of a *ΔIS4A ΔI-IIA* MN with acute Shab blockade at 22°C (top trace), 18°C (middle) and back at 22°C (bottom). **f**, quantification of spiking phenotypes as defined in Fig. 3a at different temperature between 18°C and 30°C for control MNs (top, non-shaded bars), *ΔIS4A ΔI-IIA* MNs (top, shaded), as well as Shab blockade in control MNs (bottom, non-shaded) and in *ΔIS4A ΔI-IIA* MNs (bottom, shaded).

Before, we showed that reducing VGCC isoform diversity reduces the permissive Shab range in our model (Fig. 3g), but does this observation hold across different temperatures? While increases in temperature shift the entire *g_Shab_*-dependent bifurcation diagram (Fig. 3e) upward, the size of the functional Shab range is only minimally influenced (Fig. 5c). This underscores that reduced isoform diversity has distinct effects on neuronal excitability. Whereas manipulations, like temperature increases, only shift the functional range of MNs, reducing VGCC isoform diversity influences its size and therefore impacts excitability robustness. Quantifying the interactions of Shab and temperature perturbations *in vivo* confirms that MNs with full and with reduced VGCC isoform diversity show robust excitability across moderate temperature changes (18° C and 30° C) that often occur (Fig. 5f, top). By contrast, in the context of reduced Shab levels, MNs with full VGCC isoform diversity are robust up to 26° C but start failing at 30° C. MNs with reduced VGCC isoform diversity are less robust against temperature changes and start failing at 18° C, but failure percentage increases with rising temperature. Therefore, our model predicts the interactions of Shab and temperature manipulation well, so that both theory and experiment suggest that full VGCC isoform diversity, and thus a large variance of the mean half activation voltage, safeguard excitability robustness across the temperature range that is behaviorally highly relevant to fruit flies.

In sum, it is clear that the astounding number of functions that are orchestrated by combinations of ion channels in the neuronal membrane has led to a tremendous diversification of channel isoforms. However, many highly specialized isoforms are required only in specific neuron-types, or only in specific neuronal sub-compartments. For the example of the 18 *Drosophila* Ca_v_2 splice isoforms investigated here, only a subset is critically required in presynaptic active zones for synaptic vesicle release, but others are required in other cellular compartments, and importantly, flight MNs express synaptic and non-synaptic isoforms in the soma and primary neurite that affects spike generation (Bell et al., 2025). Conceptually similar, but at increased complexity, in the human brain, the VGCC 1.2 is encoded by the psychiatric risk gene CACNA1C that encodes for 251 splice isoforms (Clark et al., 2020). It is hard to imagine regulatory mechanisms for the expression of each isoform in each neuronal compartment. Instead, our findings support the view that regulatory mechanisms must exist only to target distinct isoforms to specific parts of some neuron types, but evolution has embraced the variance that results from a less regulated expression of all other isoforms, and employs that very variance to create excitability robustness.

---

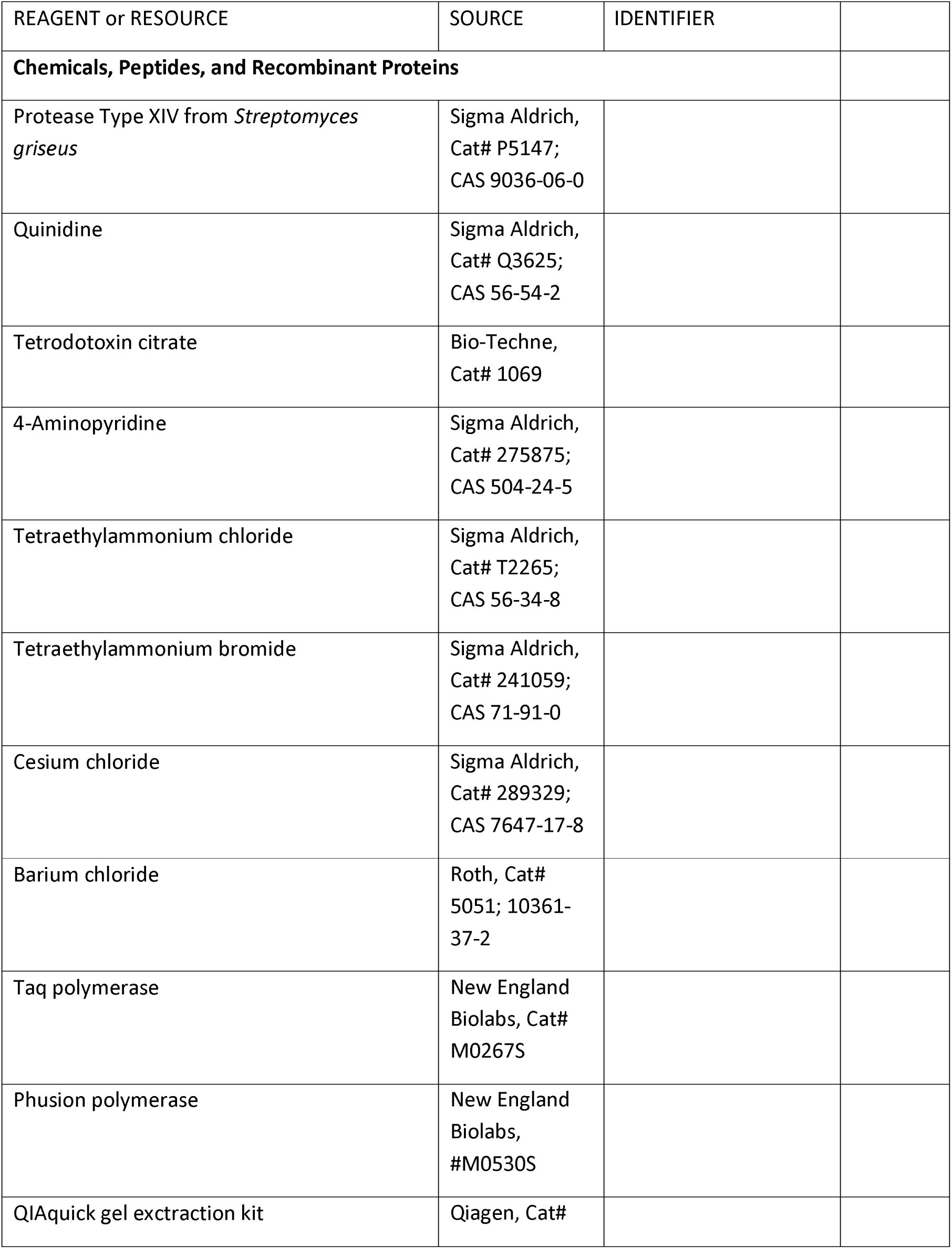
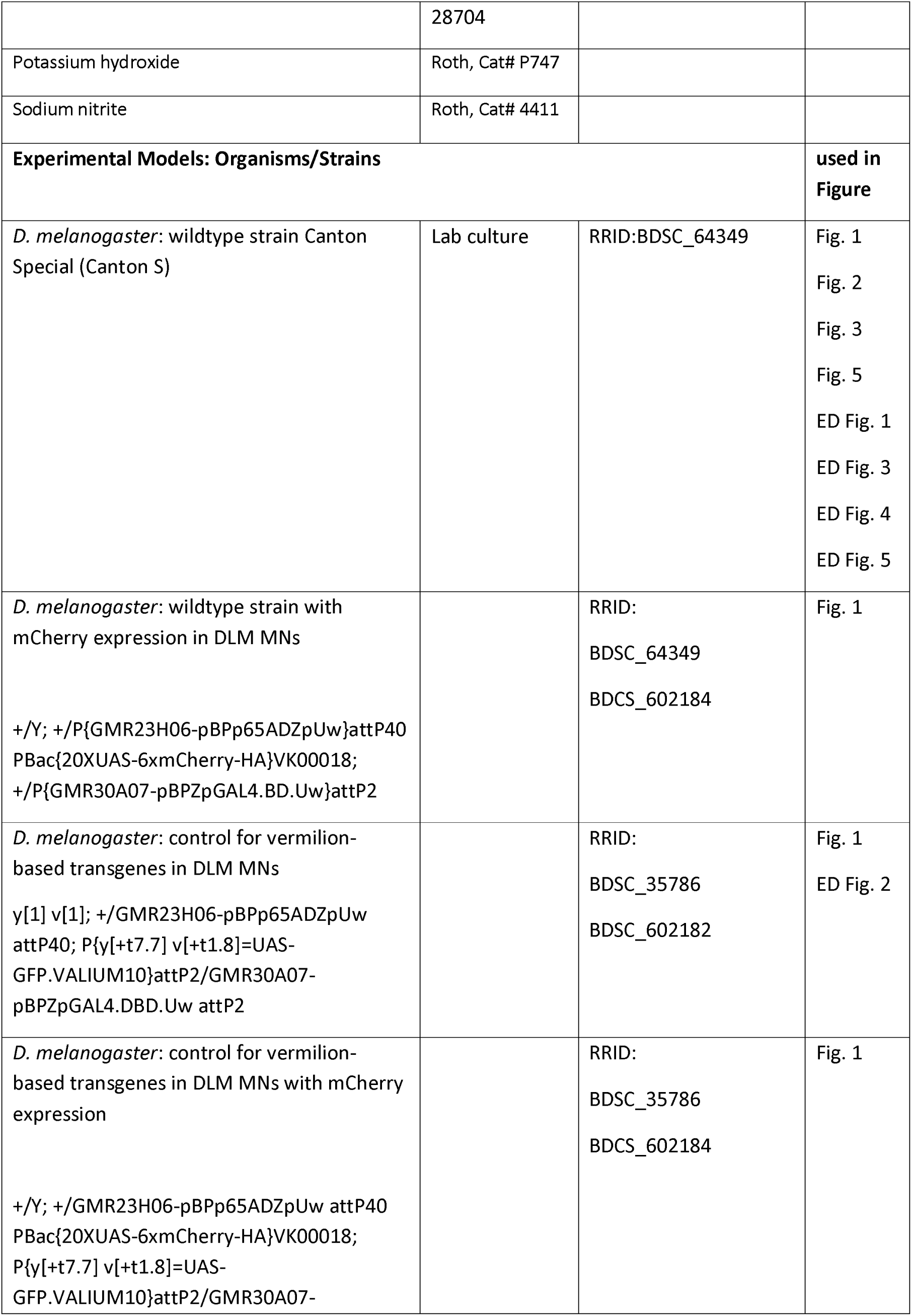
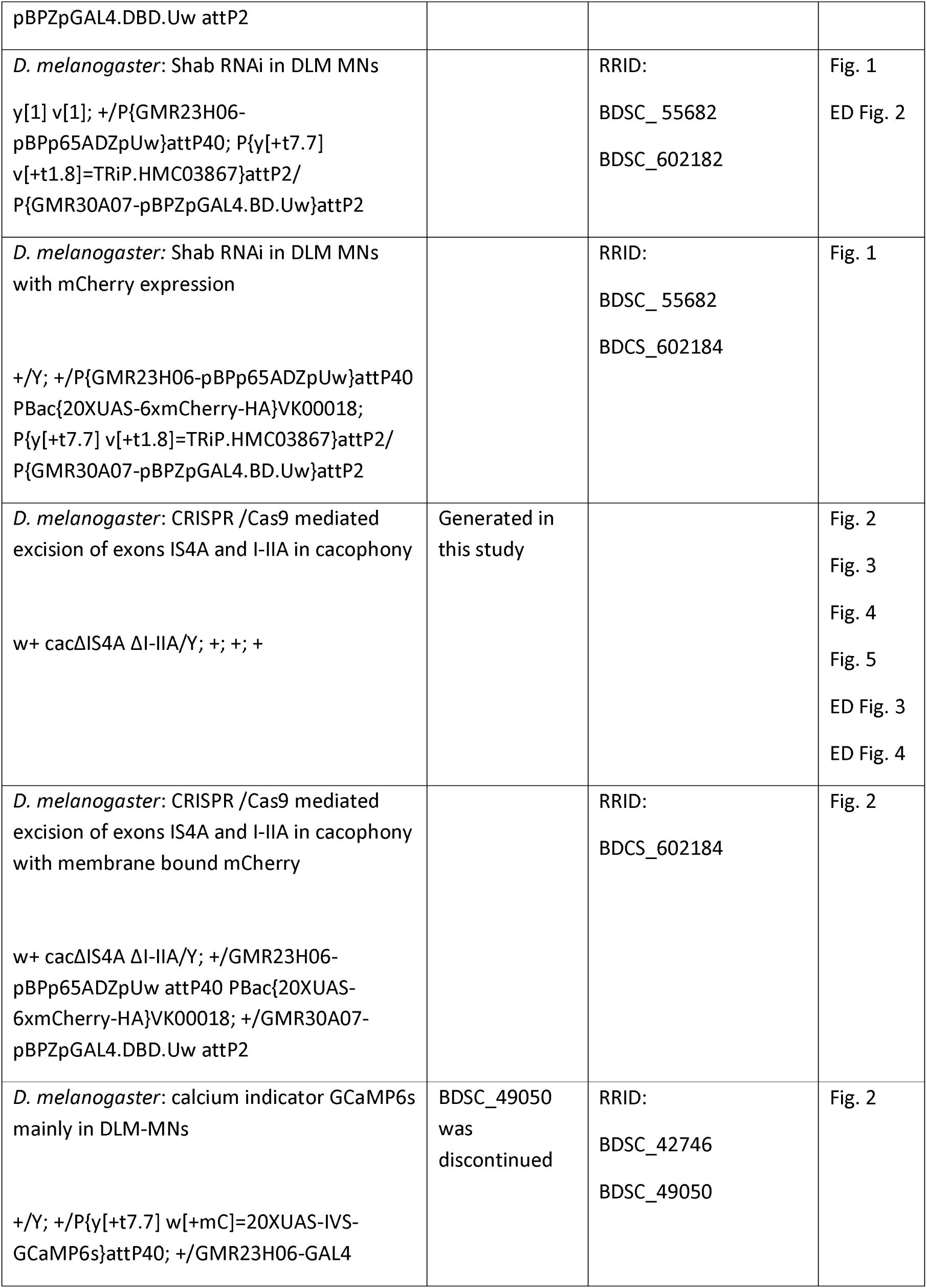
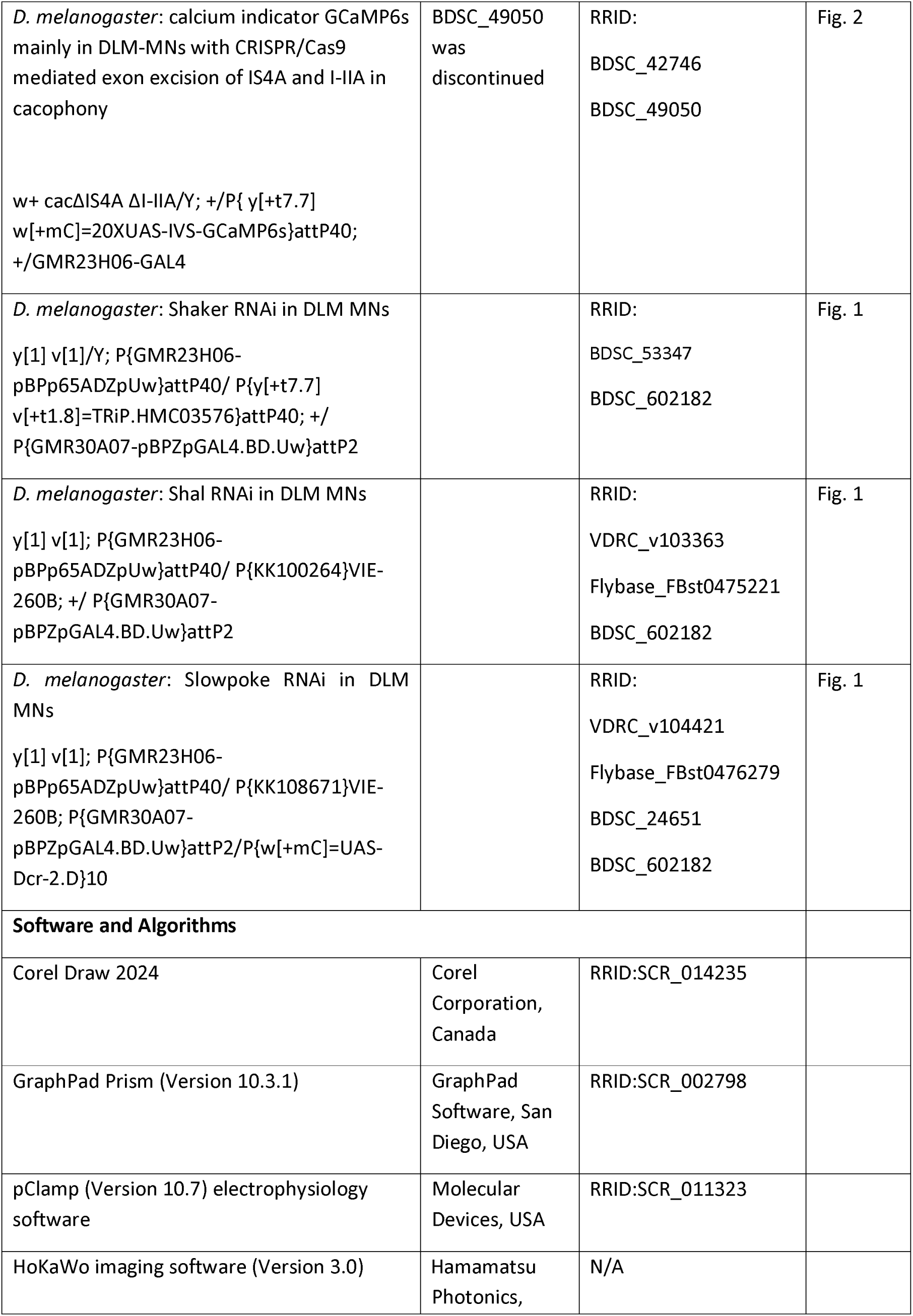
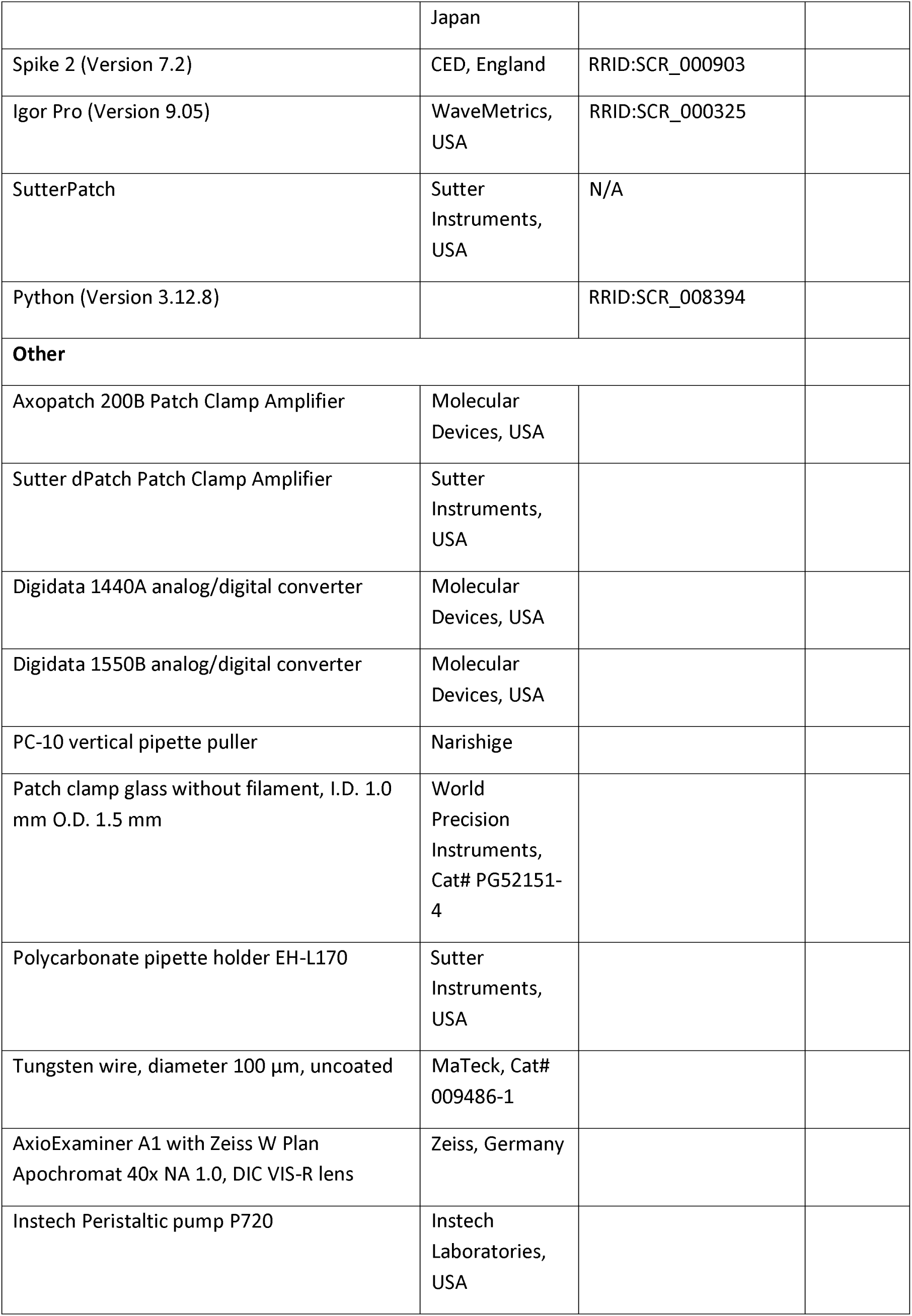
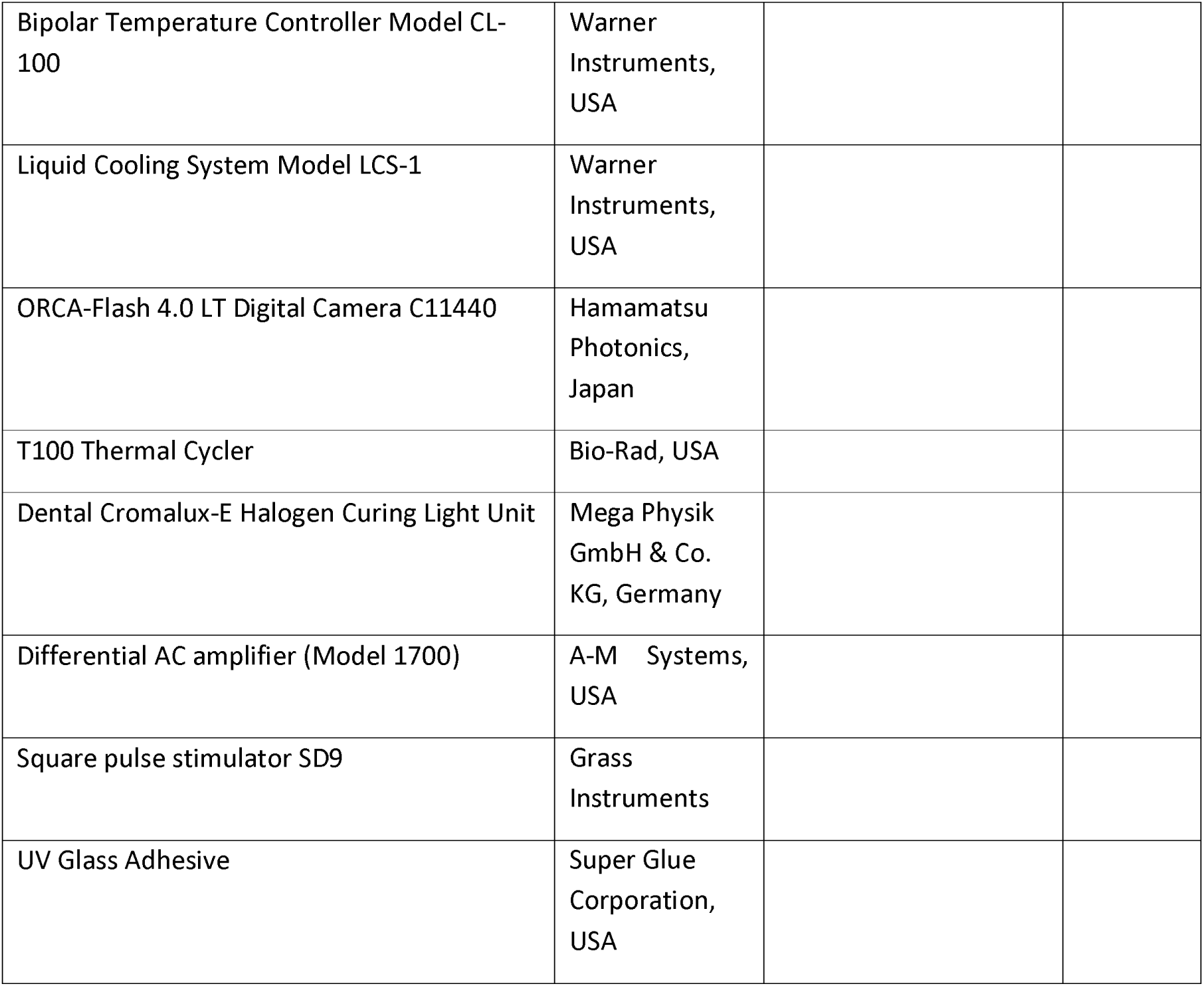
Key Resource Table

## Methods

### Animals

*Drosophila melanogaster* were reared in 68 ml plastic food vials (25 mm × 95 mm, Kisker Biotech) containing 10 ml of a standard cornmeal-based diet closed with cotton plugs. Flies were kept at 25°C and 60% humidity under a 12-hour light/dark cycle. Only male pupae at the early developmental stage P9 (as defined by Bainbridge & Bownes, 1981) or 1-2 days old adult male flies were used in experiments, except for the characterization of the delayed rectifier Shab potassium current and current clamp recordings in Shab RNAi in pupal MN5. Here, individuals of both sexes were included. A split-GAL4 driver line (BDSC# 602182), combining the driver lines GMR23H06 and GMR30A07, was used to selectively target the five DLM-MNs. The enhancer GMR23H06 drives expression of the GAL4 activation domain (AD), while the enhancer GMR30A07 drives expression of the GAL4 DNA-binding domain (DBD). Their overlapping expression patterns result in reconstituted, functional GAL4 specifically in the five DLM-MNs and two TTMs (tergorochanteral motoneurons). For fluorescent labeling of DLM-MNs membranes, mCherry-HA (BDSC# 52267) was recombined onto the second chromosome of the split-GAL4 driver line (BDSC# 602184). Only adult patch clamp recordings were performed with labeled DLM-MNs, as pupal MN5 is clearly visible by eye. Using these driver lines, a Shab knockdown TRiP transgene (BDSC# 55682), inserted at the attP2 landing site, was expressed to reduce the Shab potassium current in pupal and adult DLM-MNs. An identical genetic background, but carrying an empty VALIUM10 vector (BDSC# 35786) at the same landing site was used as a genetic control. To manipulate the number of different splice isoforms of the voltage gated calcium channel cacophony, on-locus CRISPR/Cas9 mediated exon excision was performed in CS wildtype flies to generate cac^ΔIS4A^ ^ΔI-IIA^ flies, as previously described by (Bell et al., 2024). Thus, CS flies were used as genetic control. Calcium imaging of MN5 was performed using targeted expression of GCaMP6s (BDSC# 42746) to the DLM-MNs under the control of 23H06 (BDSC# 49050, discontinued). For calcium imaging of the exon out mutants, the 23H06 driver line was recombined with UAS-GCaMP6s and then crossed into the respective exon-out genotypes or CS as genetic control. For extracellular recordings of the DLM-MN circuit, Shaker RNAi (BDSC# 53347), Shal RNAi (VDRC# v103363), and Slowpoke RNAi (VDRC# v104421) were selectively targeted to the five DLM-MNs using the split-GAL4 driver line (BDSC# 602182).

### Generation of cacophony exon-out variants

On-locus exon excision in the cacophony gene was done as described in (Bell et al., 2025) utilizing the CRISPR/Cas9 method (Doudna and Charpentier, 2014; Sternberg and Doudna, 2015). Flies carrying the Cas9 enzyme in the germline under the control of nanos (BDSC# 78781) were crossed into male flies expressing a gRNA transgene that specifically targets sequences flanking a desired exon (Table 1), which is under the control of germline U6 promoter. Single female virgins of the F1 generation were collected and back-crossed into a X-chromosomal balancer. Verification of successful exon-excision was done via PCR of the progeny. After confirmed excision of IS4A (w^+^ cac^ΔIS4A^), the procedure was repeated to excise I/IIA in the w^+^ cac^ΔIS4A^ flies for the generation of w^+^ cac^ΔIS4A^ ^ΔI-IIA^ flies. Exon excision was confirmed with a PCR (cycler program see Table 3, T100 Thermal Cycler, Bio-Rad) using primers for IS4A and I-IIA (Table 2) and Taq polymerase (New England Biolabs, #M0267S) with subsequent 1% agarose gel electrophoresis. For evaluation of break points in the confirmed double exon out flies (w^+^ cac^ΔIS4A^ ^ΔI-IIA^), a PCR with high genomic DNA yield using a Phusion polymerase (New England Biolabs, #M0530S) was employed for subsequent gel extraction and DNA purification. Extraction and purification were conducted as described in the provided manual (QIAGEN #28704). Confirmation of gene sequences flanking the excised exons was done using next generation sequencing (StarSeq GmbH, Mainz) with primers provided in Table 2.

**Table 1.**
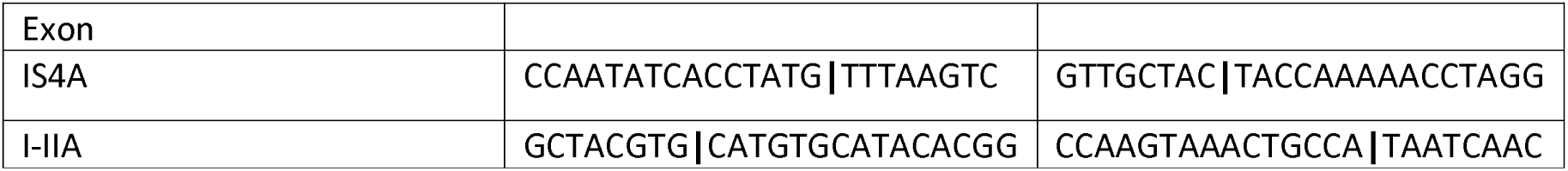
gRNA sequences. Lines depict break points

**Table 2.**
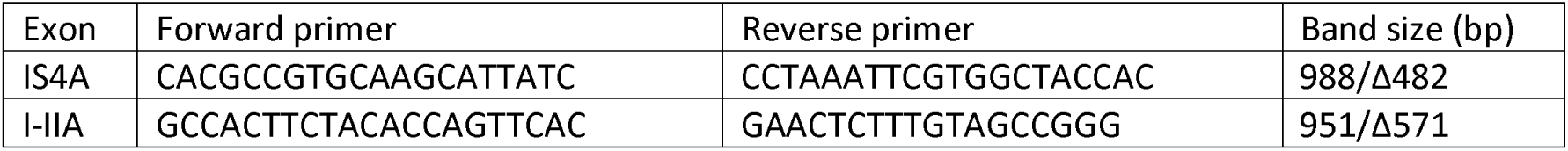
Primers for exon-out verification

**Table 3.** Cycler program for exon-out verification

| Step | Temperature |  | Duration |
| --- | --- | --- | --- |
| 1 | 95°C |  | 3:00 min |
| 2 | 95°C |  | 0:30 min |
| 3 | 57°C | -1°C per cycle | 0:45 min |
| 4 | 68°C |  | 1:30 min |
| 5 | Go to step 2 | 10x |  |
| 6 | 95°C |  | 0:30 min |
| 7 | 52°C |  | 0:45 min |
| 8 | 68°C |  | 1:30 min |
| 9 | Go to step 6 | 24x |  |
| 10 | 68°C |  | 5:00 min |
| 11 | 4°C | | ∞ |

### Pupal and adult dissection

Preparation of the dorsal ventral nerve cord (VNC) for whole-cell patch clamp recording of MN5 is performed as described by (Ryglewski & Duch, 2009). For dissection of pupal stage P9, as defined by (Bainbridge & Bownes, 1981), the puparium protecting the developing pupa is carefully removed by first cutting the operculum with an iris scissor and then unwrapping the pupa from anterior to posterior side using a forceps. Subsequently, the pupa is fixated with its dorsal side up in a ∼35 mm sylgard-coated Petri dish by using a very sharp pin placed in the very end of the abdomen before submerging it with normal saline. The hypoderm that surrounds the body becomes visible and is locally removed around the head using a pair of scissors, facilitating the following steps of dissection. A second very sharp pin is placed in the head while carefully stretching the pupa. The next steps are similar to the adult dissection of the VNC. Note that the adult dorsal preparation begins by cutting the legs and wings close to the thorax before pinning the fly with its dorsal side up in a dish. Then the cuticle of the pupa or adult fly is opened along the midline from posterior side of the abdomen to the anterior side of the thorax. Two pins are used to unfold the thorax and placed in the DLM muscles. While in the pupa the organs and loose debris that cover the VNC were carefully removed by rinsing the preparation with normal saline, the clearly visible esophagus and gut, covering the VNC of the adult fly were removed using a fine forceps. In both preparations the now visible salivary glands flanking the VNC were carefully removed and the head was cut to facilitate electrode access to the pro- and mesothoracic neuromere. Finally, the dissected fly was gently rinsed with normal saline.

### Solutions

#### Normal Saline

128 mM NaCl, 2 mM KCl, 4 mM MgCl_2_, 1.8 mM CaCl_2_, 5 mM HEPES, and ∼35 mM sucrose. Osmolality was adjusted to 300-310 mOsm kg^-1^ with sucrose. pH was adjusted to 7.24-7.26 with 1 N NaOH.

#### Normal internal solution

140 mM potassium gluconate, 2 mM MgCl_2_, 2 mM Mg-ATP, 11 mM EGTA, 10 mM HEPES, and, if needed, sucrose to adjust osmolality to 300-310 mOsm kg^-1^. pH was adjusted to 7.25-7.26 with 1 N KOH.

#### TEA saline

93.4 mM NaCl, 5 mM KCl, 4 mM MgCl_2_, 1.8 mM CaCl_2_, 30 mM TEA-Cl, 2 mM 4-AP, 1.8 mM BaCl_2_, 5 mM HEPES, and ∼35 mM sucrose. Osmolality was adjusted to 300-310 mOsm kg^-1^. pH was adjusted to 7.25-7.26 with 1 N HCl.

#### Internal TEA solution

131 mM CsCl, 0.5 mM CaCl_2_, 2 mM ATP-Mg, 5 mM EGTA, 20 mM TEA-Br, 0.5 mM 4-AP, 10 HEPES, and, if needed, sucrose to adjust osmolality to 300-310 mOsm kg^-1^. pH was adjusted to 7.25-7.26 with 1 N CsOH.

#### Calcium imaging saline

116 mM NaCl, 2 mM KCl, 4 mM MgCl_2_, 5 mM CaCl_2_, 5 mM HEPES, and ∼35 mM sucrose. Osmolality was adjusted to 300-310 mOsm kg^-1^ with sucrose. pH was adjusted to 7.24-7.26 with 1 N NaOH

#### Internal calcium imaging solution

140 mM potassium gluconate, 2 mM Mg-ATP, 2 mM MgCl_2_, 10 mM phosphocreatineditris, 0.3 mM Na_2_GTP, 10 mM HEPES and, if needed, sucrose to adjust osmolality to 300-310 mOsm kg^-1^. pH was adjusted to 7.25-7.26 with 1 N KOH.

#### *In situ* whole-cell patch clamp recording

Whole cell patch clamp recordings in pupal and adult MN5 were performed similar to (Ryglewski & Duch, 2012). The VNC preparation was positioned on a fixed stage microscope (AxioExaminer A1, Zeiss, Germany) with a 40x water dipping lens (Zeiss Objective W Plan-Apochromat 40x/1.0 DIC M27). MN5 is located at the dorsal side of the mesothoracic neuromere and surrounded by a ganglionic sheath. For successful seal formation it is mandatory to clean the soma by applying a short protease treatment. A glass electrode was pulled from borosilicate glass capillaries (O.D. 1.5 mm, I.D. 1.0 mm, World Precision Instruments) using a vertical puller (Narisighe PC-10) before breaking the tip using a forceps. The tip size was max 30% of the cell body’s diameter, ensuring a precise pressure control, and filled with 1% protease type XIV (from Streptomyces griseus, Sigma Aldrich, Cat# P5147) in normal saline. The electrode was then inserted into the electrode holder (EH-P170, Sutter Instrument) which was connected to a head stage (0.1x gain whole cell configuration, Molecular Devices, USA) on a micromanipulator system (Multi-Manipulator Controller MPC-200 and ROE-200, Sutter Instruments, USA). Positive and negative pressure was gently applied to loosen and remove other cells and debris in close proximity to the soma. Protease application was kept as short as possible to avoid damaging the cell. After cleaning the soma, the perfusion system (Instech peristaltic pump P720, Instech Laboratories, USA) with a flow rate of ∼1 ml min^-1^ was inserted into the bath to allow rigorous removal of protease. Bath volume should be minimized to allow fast exchange of solution and kept at a constant volume during recording to avoid offset changes and changes in series resistance. For patch clamp application, the enzyme pipette was replaced by a patch pipette (O.D. 1.5 mm, I.D. 1.0 mm, World Precision Instruments) filled with internal solution. Depending on the recording solutions, pipette tip resistance was between 5.5 MΩ to 6 MΩ for current clamp and potassium current recordings and between 3.5 MΩ to 4 MΩ for calcium current recordings. Before seal formation, slight positive pressure was applied to prevent influx of extracellular solution into the electrode’s tip and the offset was adjusted when approaching the soma. A stable giga seal (8.2 MΩ) was formed by gently applying negative pressure. Electrode capacitance was compensated before applying slight and short negative pressure to rapture the cell membrane to gain excess to the intracellular milieu of MN5. Whole-cell capacitance (P9: ∼130 pF, adult: ∼150 pF) and series resistance (P9: ∼10 MΩ (22°C) (current clamp and potassium currents), ∼12 MΩ (22°C) (calcium currents), adult: ∼7 MΩ (22°C) (current clamp and potassium currents)) were determined and series resistance (Rs) was compensated with a maximum correction of 60% (lag filter 10 µs) and the effect of Rs on charging the membrane was corrected using a maximal prediction of 95% depending on the maximum applied voltage step. Holding currents of less than -100 pA for current clamp recordings and potassium current measurements were considered good quality when clamped at -70 mV, whereas a holding current of max -200 pA was considered appropriate quality for voltage clamp recordings of calcium currents. Resting membrane potential in pupal and adult MN5 was ∼ -60 mV (22°C) without current injection. With these quality criteria and aforementioned cell parameters, recordings remained stable for at least 30 min. During recording, Rs was constantly monitored and recordings were discarded if Rs changed. Recording temperature was controlled via a temperature and cooling system (Bipolar Temperature Controller Model CL-100 and Liquid Cooling System Model LCS-1, Warner Instruments, USA) and unless otherwise noted, carried out at 22°C (see Temperature paradigms). Patch clamp recordings were mainly conducted using an Axopatch 200B amplifier (Molecular Devices, USA) with a sampling rate of 50 kHz (DigiData 1440A, Molecular Devices), filtered through a 5 kHz low-pass Bessel filter, and recorded with 5x gain in pCLAMP 10.7 software (Molecular Devices). For dynamic clamp, a different amplifier was used (see Dynamic clamp).

The intrinsic excitability of MN5 was determined using square pulse injections from 0 to 0.5 nA with 0.05 nA increments (1000 ms per sweep). Total potassium currents were elicited by voltage steps from -90 mV to +50 mV in 10 mV increments (500 ms per sweep) from a holding potential of -90 mV. Delayed rectifier potassium currents were elicited by voltage steps from -90 mV to +50 mV in 10 mV increments (500 ms per sweep) from a holding potential of -20 mV to inactivate the A-type potassium currents. A-type currents were calculated by subtracting the delayed rectifier currents from the total potassium currents. Calcium currents were elicited by voltage steps from -90 mV to +20 mV in 10 mV increments (200 ms per sweep) from a holding potential of -90 mV. Input resistance was calculated from the slope of the IV curve below -60 mV and subtracted off-line. Minimal input resistance in voltage clamp and current clamp recordings was ∼100 MΩ (adult) and ∼120 MΩ (P9). Maximal input resistance was ∼200 MΩ. Data was analyzed in Clampfit 10.7 (Molecular Devices).

#### Pharmacology

Measurements of total potassium currents were performed in normal saline using 100 nM TTX to block fast voltage gated sodium channels. Voltage-dependent potassium currents were measured in calcium-free normal saline with 100 nM TTX to avoid activation of calcium-dependent potassium currents. Measurements of calcium currents were performed in TEA external solution using 100 mM TTX. For pharmacological block of delayed rectifier potassium channel Shab 100 µM quinidine (C. F. Wu, personal communication) was applied 2 min before and during recording using a perfusion system with ∼1 ml min^-1^ flow rate. As quinidine is a lipophilic drug that bocks the Shab channel from the intracellular side of the pore (Gomez-Lagunas, 2010), pre-depolarization of the membrane significantly enhances Shab blockade efficiency.

### Temperature paradigms

Bath temperature was controlled as described in (*in situ* whole-cell patch clamp recording). We applied a temperature range from 18°C, 22°C, 26°C and 30°C. Temperatures were adjusted with maximal 1°C per minute. Cells were allowed to adapt 2 min after reaching target temperature (± 0.2°C). Only one cell per fly was recorded after applying different recording temperatures.

#### Dynamic clamp

Dynamic clamp (Prinz et al., 2004; Sharp et al., 1993) was carried out to acutely inject virtual ion channel conductances or to virtually block conductances in MN5. Using the dPatch amplifier from Sutter Instruments, recordings were filtered through a 10 kHz low-pass Bessel filter, sampled with 100 kHz and the calculation of the dynamic current was updated with a rate of 250 kHz. Data was recorded in the SutterPatch environment implemented in IgorPro9. The virtual conductances were simulated based on the Hodgkin-Huxley model (Hodgkin & Huxley, 1952) which describes ionic currents by gating variables that change as a function over time for a given membrane potential. The delayed rectifier potassium current is given by

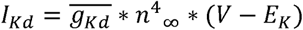

where

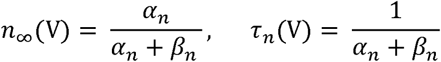

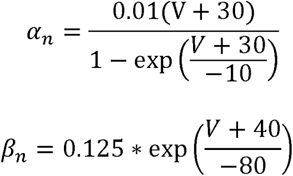

The calcium difference current (see Reduction to an effective difference current) derived in this study is given by

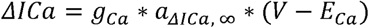

where

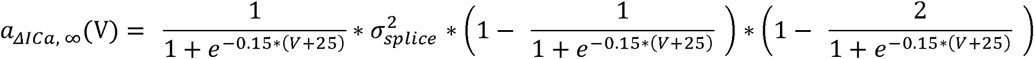

See section ‘Reduction to an effective difference current’ for its derivation.

Further parameters used are: E_K_ = -107 mV, E_Ca_ = 120 mV, *g*_Kd_ = 4 nS, g_Ca_ = 1.8 nS, and σ^2^_*splice*_ = 2. Reversal potentials were estimated based on the composition of the recording solutions. Maximal conductance value for I_Kd_ was chosen based on the maximal Shab current amplitude measured in MN5. The maximal conductance value for *ΔICα* equals the maximal amplitude of the repolarizing current component, calculated from the difference between the mean wildtype IV curve and double exon out IV curve obtained in MN5 (see Fig. 2d).

### Extracellular recording of DLM MNs

Performed as described in (Hürkey et al., 2023). Spike frequencies and phase relations of DLM-MNs 4 and 5 during tethered flight behavior were recorded extracellularly from the electrical activity of their respective target muscle fibers. MN4 exclusively innervates DLM fiber 4 and MN5 jointly innervates muscle fibers 5 and 6, allowing precise identification of individual MN spike times via the large, transient postsynaptic muscle fiber depolarization that is caused by each MN action potential.

The electromyographic recordings were prepared by cold-anesthetizing 1-2 days old male flies on ice for 10 s. They were then transferred to a metal cooling plate (3 - 6 °C) and a metal hook (0.1 mm tungsten wire clamped to a 5 cm carbon rod) was glued between head and thorax using UV Glass Adhesive (Super Glue Corporation). The glue was cured under ultraviolet light (400 - 500 nm, Mega Physik Dental Cromalux-E Halogen Curing Light Unit) for 60 s. The flies were then allowed to recover from anesthesia for 10 mins and subsequently suspended within the recording-setup to a mechanically operated micro-manipulator.

Additional micro-manipulators were used to insert the reference-electrode dorsally between the fourth and fifth abdominal segment and the recording-electrode dorsally between the left anterior dorsocentral bristle and the mid-sagittal plane through muscle fibers 5/6 into muscle fiber 4 (both electrodes were made from sharpened 0.1 mm tungsten wire, see below.). This reliably allows extracellular recording of MN5 and MN4 spike times. The earliest muscle fiber spike signals that appear upon insertion stem from muscle fibers 5/6 and the next spike signals appearing upon deeper insertion stem from muscle fiber 4. Since the signal amplitude increases with insertion depth of the electrode into each muscle fiber, large amplitude spikes can be unambiguously identified to stem from muscle 5/6 and small spikes to stem from muscle fiber 4.

Wingbeat frequency was recorded via a red laser light barrier positioned in such a way that each wing stroke interrupts the laser beam once. Using a differential AC Amplifier (Model 1700, A-M Systems) the analog muscle fiber recordings were amplified 100-fold, low-pass filtered at 100 Hz and high-pass filtered at 500 Hz before being digitized by a Digidata 1550B (Molecular Devices) at a sampling rate of 20 kHz. The signal of the laser light barrier was not amplified and digitized directly. Recordings were captured for up to 10 min using the software AxoScope v.10.7.0.3 (Molecular Devices) and saved as .abf-files. Subsequent spike identification and export of spike times to .txt-files were performed in Spike2 (v.7.2) via semi-automatic template-matching of spike-shapes and amplitudes using the built-in Wavemark function. Infrequently, MN5 and MN4 fire (nearly) simultaneously, resulting in constructive interference of their respective voltage signals within the one trace that results from recording both MNs with the same electrode. This was resolved during manual proof reading using the built-in Edit Wavemark and Split Spike function which yields two distinct, superimposed spike-shapes whose exact spike times can then be exported. Spiking frequencies and histograms of MN5 to MN4 phase relations were calculated via a custom Python (v. 3.12.8) script using the modules os and re and the packages matplotlib, numpy, pandas and scipy. In order to plot the phase-histograms, each MN4 interspike interval (ISI) within each recording was normalized to 1 (phase 0 to 1) and divided into 100 bins. The MN5 spikes within each MN4 inter spike interval (ISI) were then assigned to the bin they occurred in, and the absolute count of MN5 spikes within each MN4 bin across one recording was calculated. This count was then normalized to the total count of MN5 spikes to yield relative counts that allow further averaging of phase-histograms of all animals within one genotype.

### Electrolytic sharpening of electrodes

Recording and reference electrodes for electromyographic recordings were fashioned from 15 mm sections of 0.1 mm diameter tungsten wire crimped to circuit board pins. Each electrode was then sharpened electrolytically by repeatedly dipping its wire tip into a droplet of a NaNO2/KOH solution (10.3C⍰ NaNO2 and 6.05⍰M KOH in double-distilled H2O) while a monophasic current of 100 Hz, 40 V and 1 ms pulse-duration was applied between electrode and solution using a Grass SD9 square pulse stimulator.

### Calcium Imaging

Activity-dependent calcium influx in one of two present pupal MN5 was measured by expressing the genetically encoded calcium indicator GCaMP6s in the DLM-MN ensemble. The above described whole-cell patch clamp method was applied to elicit calcium currents by somatic current injection of 1 nA ramps (1000 ms). The change in fluorescence (ΔF/F) in the somatodendritic region and the initial axonal segment was recorded with HoKaWo imaging software (Version 3.0) using an ORCA-Flash 4.0 LT Digital Camera C11440 (Hamamatsu Photonics) installed on a fluorescence microscope (AxioExaminer A1 with Zeiss W Plan Apochromat 40x NA 1.0, DIC VIS-R lens, Zeiss, Germany). Exposure time was 75 ms. Analysis was done in HoKaWo by selecting several regions of interest (ROIs) in the primary neurite, the dendrites, and the axon and an additional ROI for background subtraction. The data frames consisting of the gray values for each frame per ROI were further analyzed in Excel (Version 2016, Microsoft) to calculate the change in fluorescence.

### Statistics

Statistical analyses were performed using GraphPad Prism (Version 10). Data were first assessed for normal distribution using the Shapiro-Wilk test. For normally distributed data, pairwise comparisons were conducted using the Student’s t-test, while comparisons across multiple groups was done employing a one-way parametric ANOVA followed by a Tukey multiple comparison post hoc test. For pairwise comparison of not normally distributed datasets the Mann-Whitney U-test was applied, while comparisons of multiple groups were performed using a one-way non-parametric Kruskal-Wollis ANOVA with Dunn’s multiple comparison post hoc test. Data are represented as mean ± standard deviation (SD) or standard error of the mean (SEM), as indicated. Statistical significance was denoted as follows: * p < 0.05, ** p < 0.01, *** p < 0.001.

### Conductance-based MN Model

MNs are described by a single-compartment conductance-based model based on (Berger & Crook, 2015). Previously (Hürkey et al., 2023), model currents were adapted to explain the mechanism behind splay state generation in the MN1-5 network. To better match our experimental recordings, all voltages (i.e., V_1/2_ and Nernst Potentials) were shifted by 10 mV. Further, we implemented temperature dependent kinetics. For network simulations, 5 identical single-neuron models were coupled by linear non-rectifying gap junction currents, *I*_gap_, following Hürkey et al., 2023. The current-balance equation of the neuron reads:

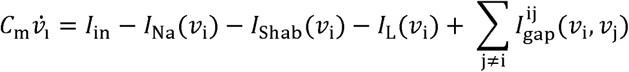

where *C_m_* is the membrane capacitance, ν the membrane voltage and *I*_in_ the input current. The sodium current *I*_Na_, potassium current *I*_Shab_, and leak current *I*_L_ are defined as:

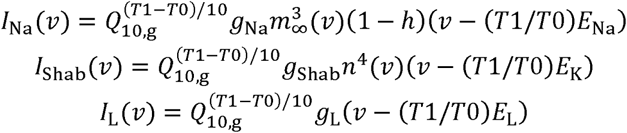

with *g*_x_ (with x ϵ Na, Shab, L) being the maximal conductance of the respective channel. With a temperature increase of 10°C, the maximal conductances are scaled by a factor of 1.3 (*Q*_10,g_ = 1.3) and the reversal potentials (*E*_x_) are shifted according to the Nernst equation. The (in-)activation gates *h*, and *n*, are defined by the following equations:

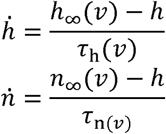

with their steady state activations (*h*_∞_, *m*_∞_, *n*_∞_) being given by:

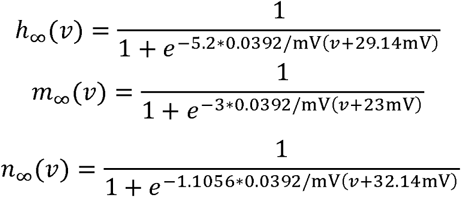

The time constants τ_h_ and τ_n_ are defined by the following equations:

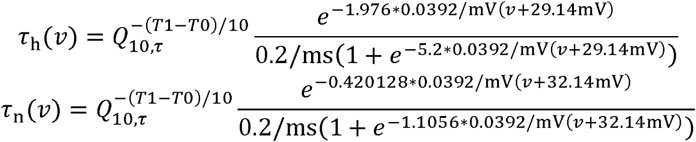

For temperature increases of 10°C, the time constants of the (in-) activation gates were scaled by a factor of 2.5 (*Q*_10,τ_ = 2.5). Gap junctional coupling is described by the following equation:

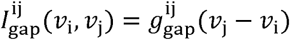

For all network simulations, a coupling strength of *g*^ij^_gap_ =43.5 pS for all *i* ≠ *j*. The splayness index was calculated as in Hürkey et al. (2023) with a perfectly splayed network having a splayness index of s=1 and a perfectly in-phase synchronized network yielding s=0. Table 4 shows the parameter values that were used throughout simulations, unless stated otherwise.

**Table 4.**
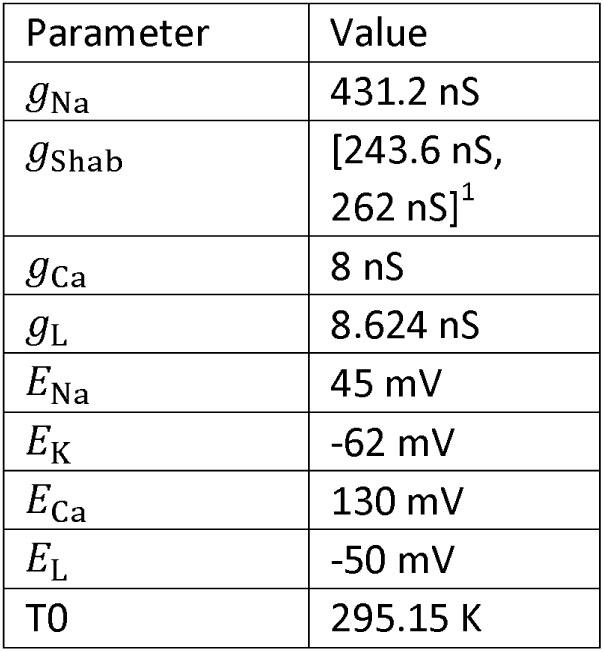
Parameter values for MN model

| Parameter | Value |
| --- | --- |
| $g_{\text{Na}}$ | 431.2 nS |
| $g_{\text{Shab}}$ | [243.6 nS, 262 nS]<sup>1</sup> |
| $g_{\text{Ca}}$ | 8 nS |
| $g_{\text{L}}$ | 8.624 nS |
| $E_{\text{Na}}$ | 45 mV |
| $E_{\text{K}}$ | -62 mV |
| $E_{\text{Ca}}$ | 130 mV |
| $E_{\text{L}}$ | -50 mV |
| T0 | 295.15 K |

<sup>1</sup> For models with 4 or 18 splice variants, respectively (see below for implementation of cacophony splice variants). $g_{\text{Shab}}$ values were chosen such that the models operated at the SNL point.

### *Cacophony* splice variant model and analysis

To account for heterogeneous splice variants of the *Cacopohony* channel, *S*=18 different sustained Ca^2+^ channels were added to the model in equal proportion (Drion et al., 2015),

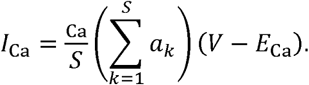

Each gating variable *a_k_* of the splice currents is assumed to follow its own kinetic equation with an activation curve that differs from the other splice isoforms only in a shift of its half-activation voltage by ⍰*_k_*,

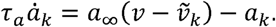

Here the sigmoidal activation curve, *a*_∞_(ν), is based on experimentally acquired data, centered at a half-activation voltage of *v_½_* = -20 mV and given by

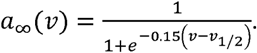

τ_*a*_ was set to 1.5 ms on the basis of our *in situ* voltage clamp recordings and equipped with temperature dependent kinetics, similar to the other channel time constants.

The ⍰*_k_* values were chosen by logit transforming 18 values evenly spaced between 0.1 and 0.9. After standardizing them to have zero mean and unit variance, the values were scaled to have a standard deviation of *σ_splice_*= 10 mV. The total maximal conductance of the calcium channel remains constant, ⍰_Ca_.

Experimentally, the number of splice isoforms of the *cacophony* channel can be reduced from 18 to 4 using the *ΔIS4A ΔI-IIA* mutant. In this case, the measured summed steady-state IV curve did not show any differences in the effective half-activation voltage or the maximal amplitude between control and *ΔIS4A ΔI-IIA*. To replicate this manipulation in the model, the four ⍰*_k_* values closest to zero were selected and their maximal conductances were set to ⍰_Ca_/4. The standard deviation of the half-activations is thereby reduced to *σ_splice_* = 1.77 mV.

### Reduction to an effective difference current

To characterize the difference current between the 18 and 4 splice variants, the current resulting from the increased splice variability is split off from the single current with half-activation at *v_½_* (*i.e.*, ⍰*_k_*=0 mV). The activation variable of a splice variant can be expressed as the difference from the average activation as *ã_k_* = *a*_k_ − *a*, such that

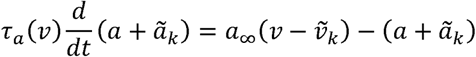

The variations, ⍰*_k_*, are drawn from a zero-mean distribution with a small standard deviation, denoted *σ*_splice_.

Since the difference between splice variants is assumed to be a shift in the activation curve by ⍰*_k_*, they can be described by the shift operator 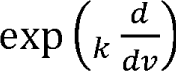. Hence the different activation curves are expressed as

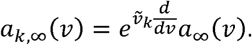

For small ⍰*_k_*, this is approximated as

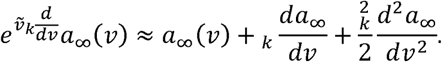

Using this approximation, one identifies the equation for the mean gating particle as

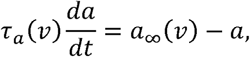

and the deviation of the different splice variants

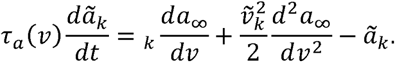

For the sake of analysis take the continuum limit of many splice variants and assume a symmetric, zero mean distribution with standard deviation *σ_splice_*, *e.g.* a Gaussian, for ⍰*_k_*. Averaging the above equation over ⍰*_k_*, the linear term in the expansion of the shift operator will cancel out and one is left with

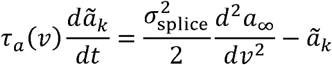

The effective activation curve of the difference current can be identified as

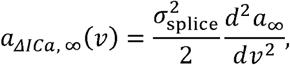

and is used in the dynamic clamp experiments.

### Software and Simulations

Model simulations were performed with the brian2 package (version 2.5.1) for python (Stimberg et al., 2019) using the Heun integrator with a time step of 5 µs. To simulate the plateau potentials (Fig. Model simulations were performed with the brian2 package (version 2.5.1) for python (Stimberg et 4a, inlets), colored noise was incorporated into the model following an Ornstein-Uhlenbeck (OU) process. The temporal evolution of the noisy input *I*_in_ was defined by the following stochastic differential equation:

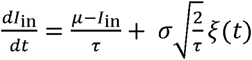

where ̃ = 20 pA is the mean input current, τ = 30 ms is the noise time constant, σ = 100 pA is the noise amplitude and ξ(*t*) is a zero-mean white noise current (using the brian2 xi variable). For the deterministic network simulations (Extended Data Fig. 6), the Runge-Kutta method (RK4) was used with a time step of 5 µs. Membrane voltages of the neurons were randomly initialized and the same initial conditions were used for each *g*_Shab_ value and for models with full and reduced VGCC isoform diversity.

The bifurcation analysis was carried out using AUTO-07P (Doedel et al., 2007).

### Onset bifurcations explain diverse spiking mechanism and network states

Neurons can exhibit different mechanisms of action potential generation that can be explained by the underlying bifurcation landscape. The bifurcation that a neuron undergoes from resting to spiking (i.e., onset bifurcation) will have consequences for different dynamical properties, like e.g., the network state. Codim-2 bifurcation diagrams highlight how different parameter values are associated with which onset bifurcation and under which conditions excitability switches take place. The bifurcation diagram in Fig. 4a shows that depending on the Shab channel conductance *g*_Shab_, associated with which onset bifurcation and under which conditions excitability switches take place. different onset bifurcations occur. For high values of *g*_Shab_, spiking is initiated through a Saddle-Node on Invariant Circle (SNIC) bifurcation. In this parameter regime, the neuron would synchronize when coupled in a network (pink region in Extended Data Fig. 5). The lower end of the SNIC region is defined by the small Saddle-Node-Loop bifurcation (sSNL, blue circle in Fig. 3e). For simplicity, when using the term SNL, we will refer to the sSNL. For *g*_Shab_ values below the SNL point, spiking commences through a small homoclinic loop (HOM, green line Fig. 3e). When connected in a network, neurons close to the SNL will show splayed out firing, which will also extend within the HOM region (green region, Extended Data Fig. 5). When further decreasing *g*_Shab_, a Hopf bifurcation defines the lower end of the spiking region as the spiking limit cycle transitions into a stable spiral and the neuron enters the depolarization block.

Within the zero-frequency limit (i.e., exactly at spiking onset), transitions from synchronous to splay states around the SNL point are sharp and well-defined. However, when moving away from this limit, the boundaries between excitability classes become blurred. This effect can be observed in Extended Data Figure 5, where high network splayness usually associated with a homoclinic spike onset extends into the SNIC region. Similarly, within the HOM region, splayness decreases to around 0.5. In the simulations, for each *g*_Shab_, the input was adjusted to be just above the saddle-node (SN) bifurcation (blue line Fig. 4a, I_in_ = I_SN_ + 1 pA). For neurons classified as homoclinic, this places the system relatively far away from the zero-frequency limit and already closer to spiking offset (i.e., depolarization block, Hopf bifurcation), resulting in higher firing rates and mixing of homoclinic and Hopf excitability features. These mixed properties can result in cluster states, where subsets of neurons synchronize and others remain desynchronized, thus reducing overall network splayness.

### Sensitivity analysis

Simulating the *ΔIS4A ΔI-IIA* mutation in the model by reducing the number of Calcium channels from 18 to 4 reduced the range of viable *Shab* levels, see Fig. 4d. To determine how sensitive this effect is to different parameter settings, we systematically varied model parameters and recorded the effect on the bifurcation diagram, specifically on the distance between the SNL point and depolarization block. While one parameter was varied, others were kept at a fixed value (for exact values, see caption Extended Data Fig. 6). The sensitivity analysis was performed for the parameters *τ_Ca_*, *lll_Ca_*, *σ*_splice_, *v*_½_.

1. When slowing down Calcium channels by increasing *τ_Ca_* (Extended Data Fig. 6a), the *Shab* range for models with full and reduced VGCC isoform diversity increases, however, the difference in their range only increases minimally with higher *τ_Ca_*. Increasing the time constants slows down the opening and closing of calcium channels during an action potential which decreases the amplitude of the total calcium current elicited from a spike but prolongs its duration (Extended Data Fig. 6b). While reducing the total amplitude of the calcium current decreases the difference in range (also see effects of *lll_Ca_* on the *Shab* range), prolonging its duration increases it. Thereby, these two effects might mitigate each other, leaving the permissive *Shab* range almost unchanged when varying *τ_Ca_*.
2. Increasing the maximal conductance of the calcium channel, *lll_Ca_*, increases the difference in *Shab* range between models with full and reduced VGCC isoform diversity (Extended Data Fig. 6c). As the overall calcium current is increased with larger *lll_Ca_* (Extended Data Fig. 6d), the difference in the dynamic and steady-state calcium current between models with a high and low variance of ⍰*_k_* also increases, which thus amplifies the difference in *Shab* range.
3. *σ_splice_* defines the variance of *v_1/2_* assuming that the different splice variants of the calcium channel vary in their half-activation (Extended Data Fig. 6e). Increasing the variance of the distribution of *v_1/2_* first increases the difference in *Shab* range up until *σ_splice_*= 20 mV. At higher *σ_splice_* values, the difference in *Shab* range gets smaller again. Increasing *σ_splice_* makes the steady-state I-V curves shallower, meaning that calcium channels would already open at very low voltages (< -80 mV for *σ_splice_* > 20 mV, Extended Data Fig. 6f). This does not match the experimentally observed activation voltages anymore (−60 mV to -50 mV, Fig. 2d) and we therefore concluded that these *σ_splice_*values are not biologically plausible anymore. Within the range that we still deemed biologically plausible (*σ_splice_* < 20 mV), the effect of increasing VGCC isoform diversity on the difference in *Shab* range is monotonic and increasing with in-creased *σ_splice_*.
4. *v_½_* also shows non-monotonic effects on the *Shab* range (Extended Data Fig. 6g). At *v_½_* = -30 mV, the reduced depolarization of the calcium current with increased VGCC isoform diversity will cover more of the voltages covered during an action potential (Extended Data Fig. 6h) and therefore move the SNL and Hopf bifurcation down. However, as most of the increased depolarization happens at higher voltages, the location of the depolarization block (i.e., Hopf line) is affected more. At -20 mV, the effectively more depolarizing and hyperpolarizing parts of the calcium current with increased VGCC isoform diversity are located such that they differentially impact action potential downstroke and refractory period (see main text for explanation). The reduced depolarization during AP downstroke moves the Hopf bifurcation down, while the increased depolarization during the refractory period moves the SNL up within the bifurcation diagram, thus increasing the functional *Shab* range. Lastly, for *v_½_* = -10 mV, calcium channels open only at higher voltages and only the more depolarizing parts of the calcium current with more VGCC isoform diversity will be covered during an AP, therefore moving the SNL and Hopf line almost equally up within the bifurcation diagram. Therefore, the difference in *Shab* range between models with full and reduced VGCC isoform diversity decreases with *v_½_* = -10 mV.

## Author contributions

S.H. conducted all in situ patch clamp recordings. L.H. and S.Hü. conducted the flight recordings. H.S. and N.N. conducted the mathematical modelling. S.H. analyzed all in situ patch clamp recordings. L.H. and S.Hü. analyzed the flight recordings. H.S., N.N., J.-H.S. analyzed the mathematical modelling.

S.R., C.D., S.H., H.S., N.N., J.-H.S., and S.S. conceptualized the work.

S.R. and C.D. supervised the experimental work, J.-H.S. and S.S. supervised the theoretical work.

S.R., C.D., S.H., H.S., J.-H.S. wrote the first draft of the manuscript.

All authors edited text and figures.

Funding by the DFG within the Research Unit FOR5289 to SS (SCHR 1239/5-1), CD (DU331/15-1), SR (RY117/4-1) and by the European Research Council (ERC) under the European Union’s Horizon 2020 research and innovation program to SS (grant agreement no. 864243)

