## Extended data figures and legends for "Robustness through variability: ion channel isoform diversity safeguards neuronal excitability": Extended Data Figures.docx

Extended Data Figures 1-6:

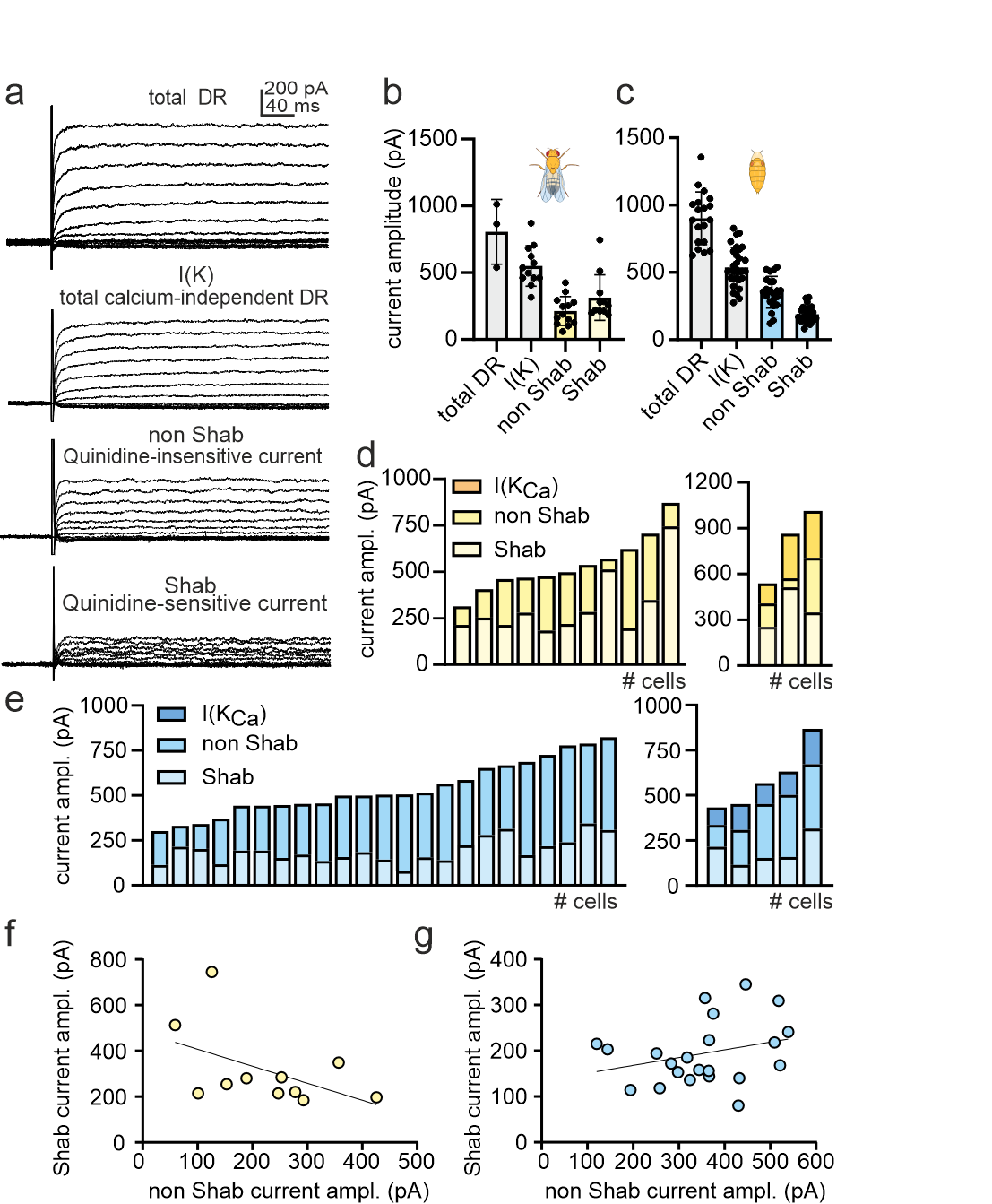

**Extended Data Figure 1.** *Variable delayed rectifier potassium current amplitudes in MN1-5.*

**a**, Representative voltage clamp traces of total delayed rectifier (DR) (top), total calcium-independent DR (I(K)) (second from top), quinidine-insensitive non-Shab (second from bottom) and quinidine-sensitive Shab current (bottom). Quantification of relative DR amplitudes in **b**, adult (total DR n = 3, I(K) n = 12, non-Shab n = 13, Shab n = 11) and **c**, developing (total DR n = 19, I(K) n = 30, non-Shab n = 22, Shab n = 22) MNs. Shab current increases during development from ~30 to 50% of total DR. Quantification of DR current relations in **d**, adult (yellow, n = 14) and **e**, developing (blue, n = 28) MNs show no clear correlations between different DR currents, but high variability in maximal current amplitudes across animals (**f**, adult Shab vs. non-Shab current, Pearson correlation r = -0.49, 95% confidence interval -0.84 to 0.16, r^2^ = 0.24, p = 0.13; g, pupal Shab vs. non-Shab current, Pearson correlation r = 0.29, 95% confidence interval -0.15 to 0.63, r^2^ = 0.08, p = 0.2). For **b** and **c** mean and SEM are shown.

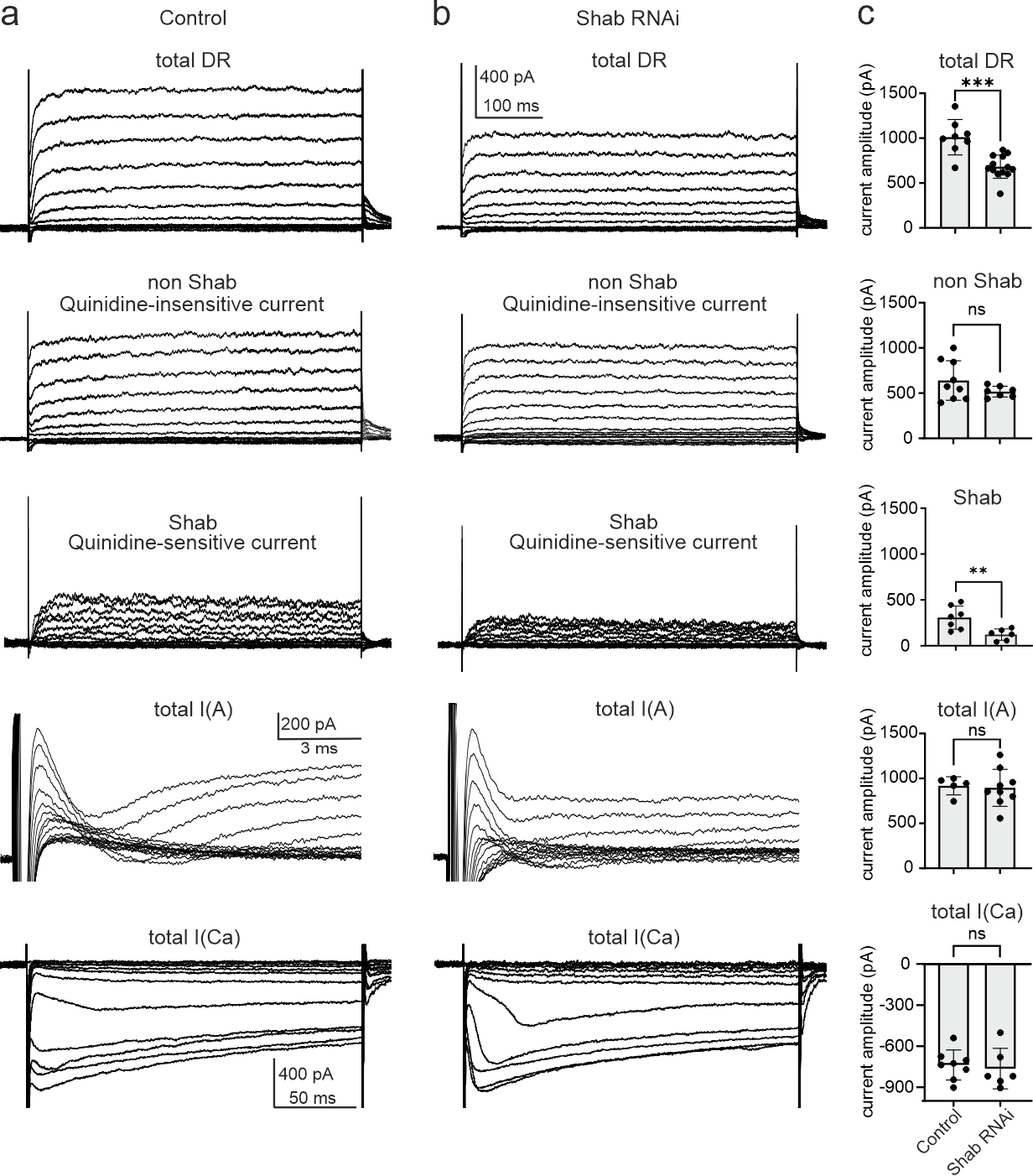

**Extended Data Figure 2.** *Potassium and calcium currents in genetic Shab manipulation are not changed.* Recordings done in pupae. Representative voltage clamp traces of total DR (top), quinidine-insensitive non-Shab (second from top), quinidine-sensitive Shab (third from top), total A-type (second from bottom) and total calcium currents (bottom) in **a**, control and **b**, Shab RNAi targeted to DLM-MNs. **c**, Quantification of maximal current amplitudes (total DR: control n = 8, Shab RNAi n = 13); non-Shab: control n = 9, Shab RNAi n = 7; Shab: control n = 7, Shab RNAi n = 6; total A-type: control n = 5, Shab RNAi n = 9; total calcium current: control n = 8, Shab RNAi n = 6) show that upon significantly reduced Shab current in Shab RNAi, mean current amplitude of the total DR is significantly reduced, whereas the remaining mean potassium and calcium current amplitudes are unchanged (unpaired t-test, total DR p = 0.0002, non-Shab p = 0.1715, Shab p = 0.0079, total A-type p = 0.8277, total calcium current p = 0.7066. Data shows mean ± SEM.

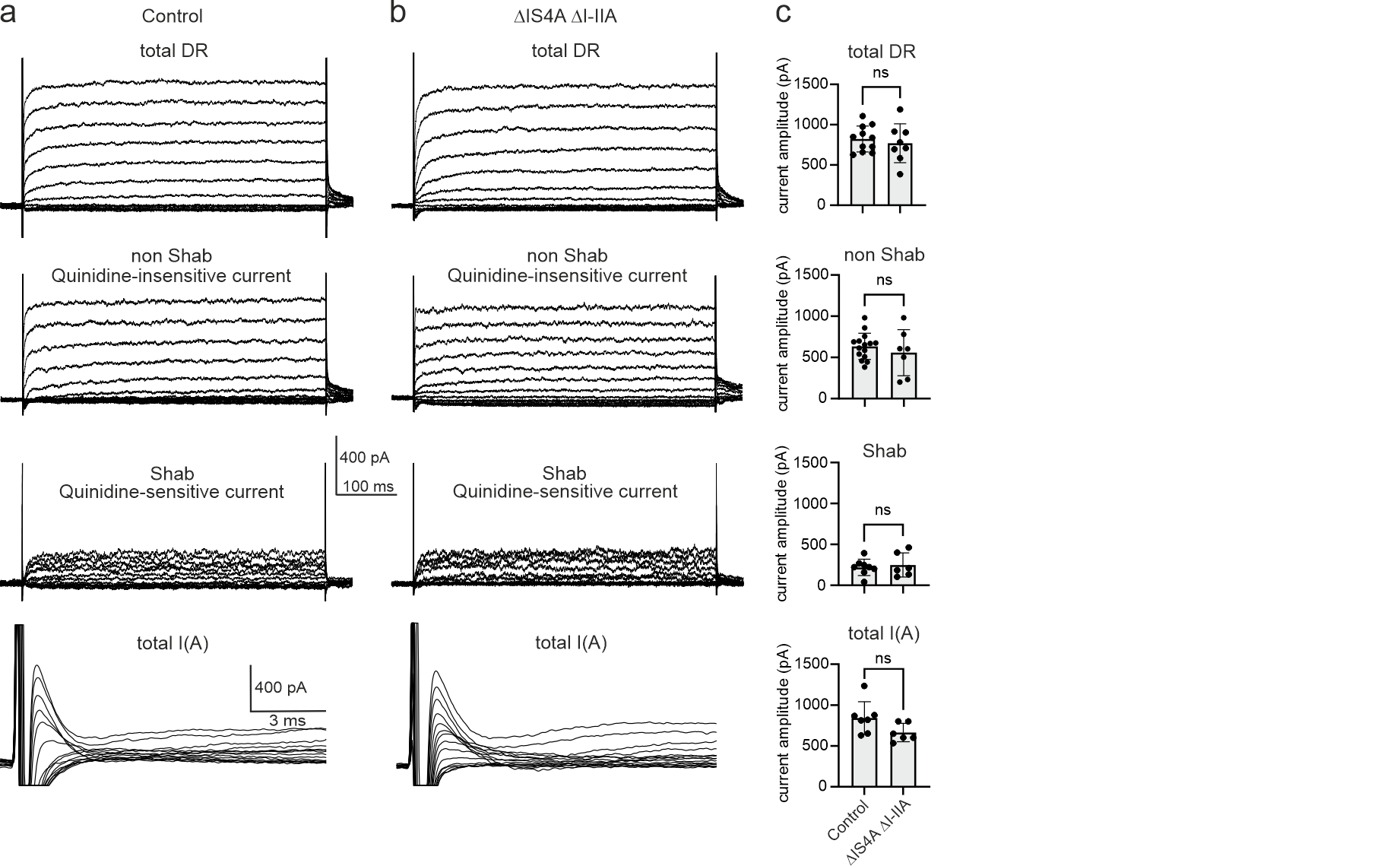

**Extended Data Figure 3.** *Potassium currents in a genetic background with reduced VGCC isoform diversity.* Recordings done in pupae. Representative voltage clamp traces of total DR (top), quinidine-insensitive non-Shab (second from top), quinidine-sensitive Shab (second from bottom), total A-type (bottom) in **a**, control and **b**, ∆IS4A ∆I-IIA. Quantification of maximal mean current amplitudes of potassium currents (total DR: control n = 11, ∆IS4A ∆I-IIA n = 8; non-Shab: control n = 15, ∆IS4A ∆I-IIA n = 7; Shab: control n = 8, ∆IS4A ∆I-IIA n = 6; total A-type: control n = 7, ∆IS4A ∆I-IIA n = 6) shows that they are not affected upon reduced VGCC isoform diversity (unpaired t-test, total DR p = 0.5754, non-Shab p = 0.4227, Shab p = 0.6618, total A-type p = 0.0796. Data shows mean ± SD.

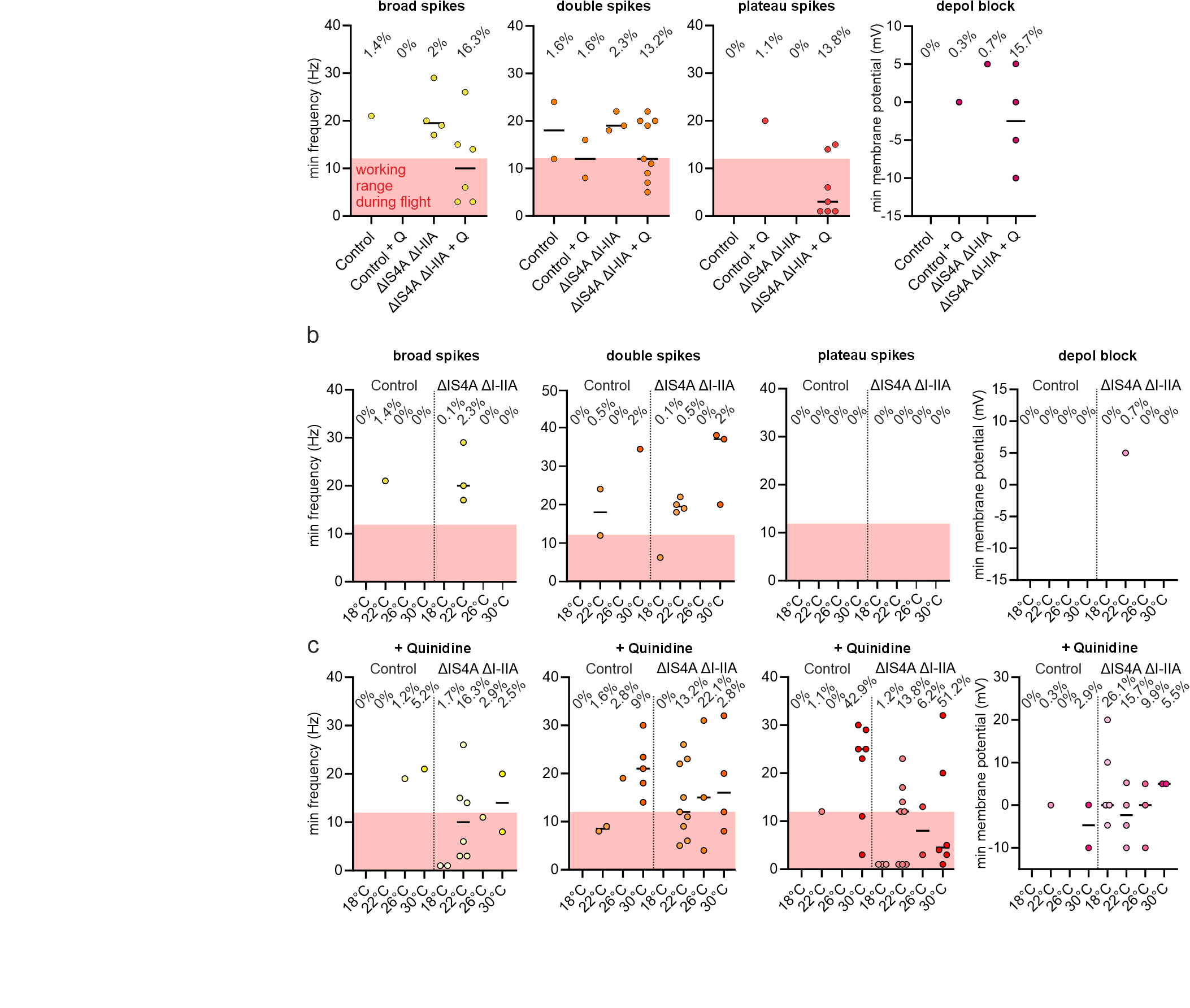

**Extended Data Figure 4.** *Quantification of spiking phenotypes within relevant frequency range during flight.* To show the *in vivo* relevance of effects summarized in figure 3c, we analyzed whether abnormal spike shapes (Fig. 3a) occur within the normal working range of MN firing during flight (3 – 12 Hz, red shaded areas. For each genotype, we quantified the minimal firing frequencies and membrane potential where different aberrant spiking phenotypes shapes start to occur. Dots represent single MNs. Numbers at the top display relative amount of time spent in a given spiking category. Red area indicates working range during flight. Reduced VGCC isoform diversity renders excitability less robust to Shab perturbation, especially in the behaviorally relevant frequency range.

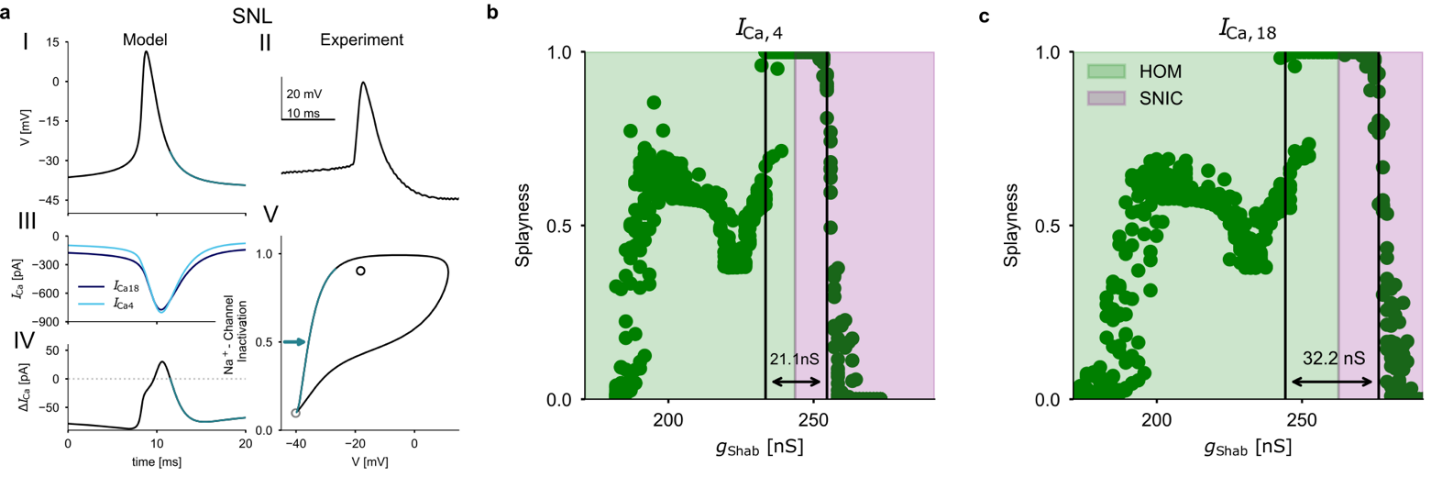

**Extended Data Figure 5.** *VGCC isoform diversity increases the Shab range permissive for splayed-out firing.* **a**, Simulations of membrane voltage and calcium current during an action potential close to the SNL point. First row shows model voltage traces during an action potential (**I**) that are similar to those recorded *in vivo* (**II**). Second row depicts the dynamic calcium currents during an action potential for models with 4 (light blue) and 18 (dark blue) VGCC isoforms (**III**). In the third row (**IV**), the differences of the dynamic currents (dynamic $\Delta I_{\mathrm{Ca}}$) are shown over the time course of the action potential. **V**, phase portrait of an action potential. The ghost of the saddle-node (i.e., old resting state) is shown as a grey open circle. An unstable spiral (i.e., the location of the depolarization block) is indicated by a black open circle. The additional depolarization through $\Delta I_{\mathrm{Ca}}$ at the end of the AP (teal lines) stabilizes the homoclinic spike onset for models with full isoform diversity. Splayness index (see methods and Hürkey et al., 2023) in models of the full MN1-5 network across different values of $g_{\mathrm{Shab}}$ (10 second simulations with 10 runs per condition) for models with reduced (*Ι*_Ca, 4_, **b**) and full isoform diversity (*Ι*_Ca, 18_, **c**). For each $g_{\mathrm{Shab}}$, input was adjusted to be just above threshold (I_in_ = I_SN_ + 1 pA). HOM (green) and SNIC (pink) excitability types are separated by the SNL bifurcation, where splayness peaks. Black vertical lines indicate the region, in which mean splayness > 0.75 across the 10 simulations. This region is increased with full isoform diversity.

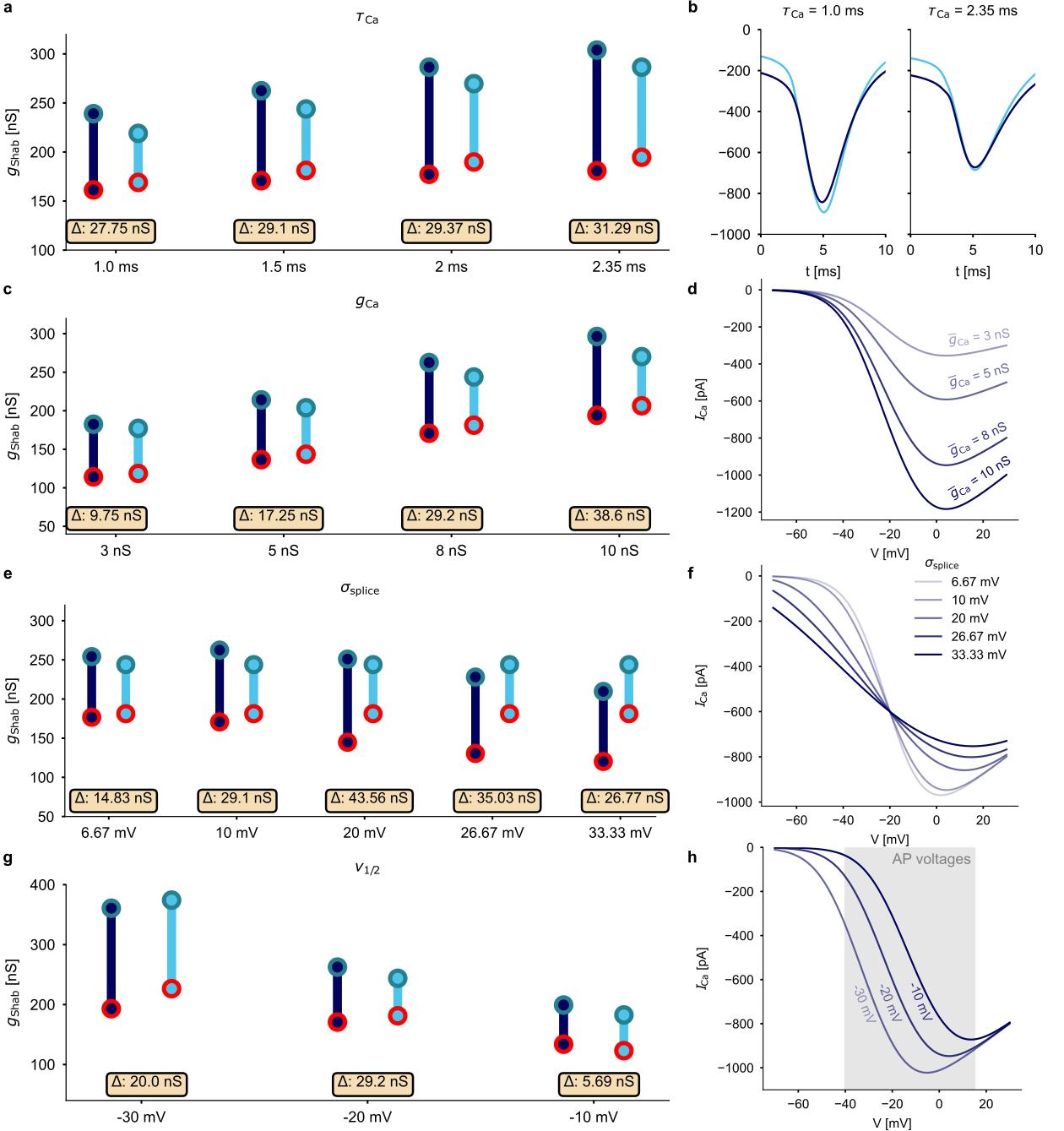

**Extended Data Figure 6.** *Sensitivity of permissive Shab range to different model parameters.* **a**, Effects on *Shab* range when increasing $\tau_{\mathrm{Ca}}$ between models with a full (N = 18, dark blue) and reduced (N = 4, light blue) isoform diversity. Teal circles show the $g_{\mathrm{Shab}}$ value of the SNL point and red circles the $g_{\mathrm{Shab}}$ value of the Hopf bifurcation. Boxes at the bottom indicate the difference in *Shab* range between the two models. **b**, Calcium currents recorded during a spike for $\tau_{\mathrm{Ca}}$ = 1ms (left) and $\tau_{\mathrm{Ca}}$ = 2.35ms (right). **c**, Effects on *Shab* range when increasing $g_{\mathrm{Ca}}$. **d**, Steady-state I-V curves for the calcium current when varying $g_{\mathrm{Ca}}$. **e**, Effects on *Shab* range when increasing $\sigma_{\mathrm{splice}}$. **f**, Steady-state I-V curves for the calcium current when varying $\sigma_{\mathrm{splice}}$. **g**, Effects on *Shab* range when varying $v_{1/2}$. **h**, Steady-state I-V curves for the calcium current when varying $v_{1/2}$. Grey area shows the voltages that are covered during an action potential (AP). For all simulations, while one of the parameters was varied, the others were fixed at $\tau_{\mathrm{Ca}}$ = 1.5ms, $g_{\mathrm{Ca}}$ = 8nS, $\sigma_{\mathrm{splice}}$ = 10 mV, and $v_{1/2}$ = -20 mV.
